# Self-organized tiling generates tissue-scale hyperuniformity during development

**DOI:** 10.64898/2026.04.30.721955

**Authors:** Sandra Siegert, Lida Kanari, Mehmet Can Uçar

## Abstract

Biological tissues require branched cellular architectures to maximize spatial coverage while minimizing redundancy. Yet, how cells decode local spatial information to collectively tile territories without a global blueprint remains a key open question. Here, we develop a biophysical theory of interacting branched cells, and show that coupling their growth to short-range repulsion drives efficient tiling with minimal territorial overlap. Our model predicts that the same local mechanism simultaneously suppresses long-range density fluctuations, driving the cellular collective toward hyperuniformity. We confirm these theoretical predictions with experiments on microglial patterning in the developing retina, and show that perturbations resulting in limited cell growth disrupt both tiling and fluctuation suppression. Our results reveal that two seemingly distinct optimization principles of biological patterning, large-scale regularity and efficient tiling, are intimately linked and can arise from a single growth-repulsion feedback, suggesting a general principle for self-organized tissue coverage.

## INTRODUCTION

Efficient spatial coverage is a recurring constraint in biological organization. Branched structures ranging from individual arborized cells to multicellular networks must optimize coverage to facilitate transport of nutrients and waste materials, sensing of environmental cues, or the surveillance of surrounding tissue while at the same time limiting redundant overlap and excessive growth [1–8]. This coverage generally does not arise from pre-patterned templates, but through growth and rearrangement governed by local feedback [9–11]. How such feedback gives rise to robust large-scale order without centralized control is therefore a fundamental question.

Cellular tiling in the developing nervous system provides a key example of this challenge [12]. Branched cells partition space to minimize overlap while maintaining uniform coverage, such as neurons [13, 14] and glial cells [12, 15–17] in the brain. At the single-cell scale, molecular and cellular mechanisms underlying these architectures are increasingly well characterized, including contact-mediated branch retraction [18–25], chemical and mechanical guidance cues [26–29], and repulsion between neighboring cells [30, 31]. Recent work also started to explore the interplay between stochastic and deterministic rules that shape individual cell morphologies [9, 23, 32]. However, connecting these mechanisms to the collective patterning of an entire macroscopic tissue remains a multiscale problem that extends beyond molecular-scale descriptions [33].

Here, we address this question by developing a theoretical framework for the collective patterning of interacting branched cells. We show that coupling cell growth to short-range intercellular repulsion drives efficient tiling of space and large-scale suppression of density fluctuations. The latter is a hallmark of hyperuniformity [34], a form of hidden long-range order previously documented primarily in static biological patterns [35, 36]. We test our framework on selforganized patterning of microglia, which collectively colonize retinal layers during postnatal development to monitor the retinal environment [37, 38]. Using complementary geometric, statistical, and topological descriptors, we show that microglia transition from an irregular early arrangement toward a uniformly tiled state with increasingly suppressed large-scale fluctuations, and that perturbing microglial morphogenesis disrupts both tiling efficiency and fluctuation suppression. To identify the physical conditions underlying this transition, we develop a coarse-grained description of growing cells with soft-core interactions, which reveals an optimal regime in the growth-repulsion plane that couples maximal tiling efficiency to hyperuniformity. These results indicate that hyperuniform organization can arise as the macroscopic consequence of a local biological requirement: efficient territorial tiling by growing cells. Together, our work establishes local growth-repulsion feedback as a general mechanism for self-organized tissue patterning.

## RESULTS

### Stochastic branching and annihilation model reproduces individual microglial morphogenesis

The retina offers a spatially extended laminar domain in which microglia colonize and establish a regular pattern across the tissue [39] (**Fig. 1a**). During development, microglia infiltrate the outer plexiform layer (OPL) from postnatal day 7 (P7) onward, and form branched arbors confined to a quasi-two-dimensional sheet of ∼4–5 *µ*m thickness across stages [40], making this tissue a well-controlled system for studying spatial patterning. To determine microglial patterning during OPL development, we reconstructed microglial morphologies in *Cx3cr1* ^GFP/−^ mice across stages P7–P30 (see **Methods**), spanning the period from colonization to functional retinal maturity [41, 42].

**FIG. 1.**
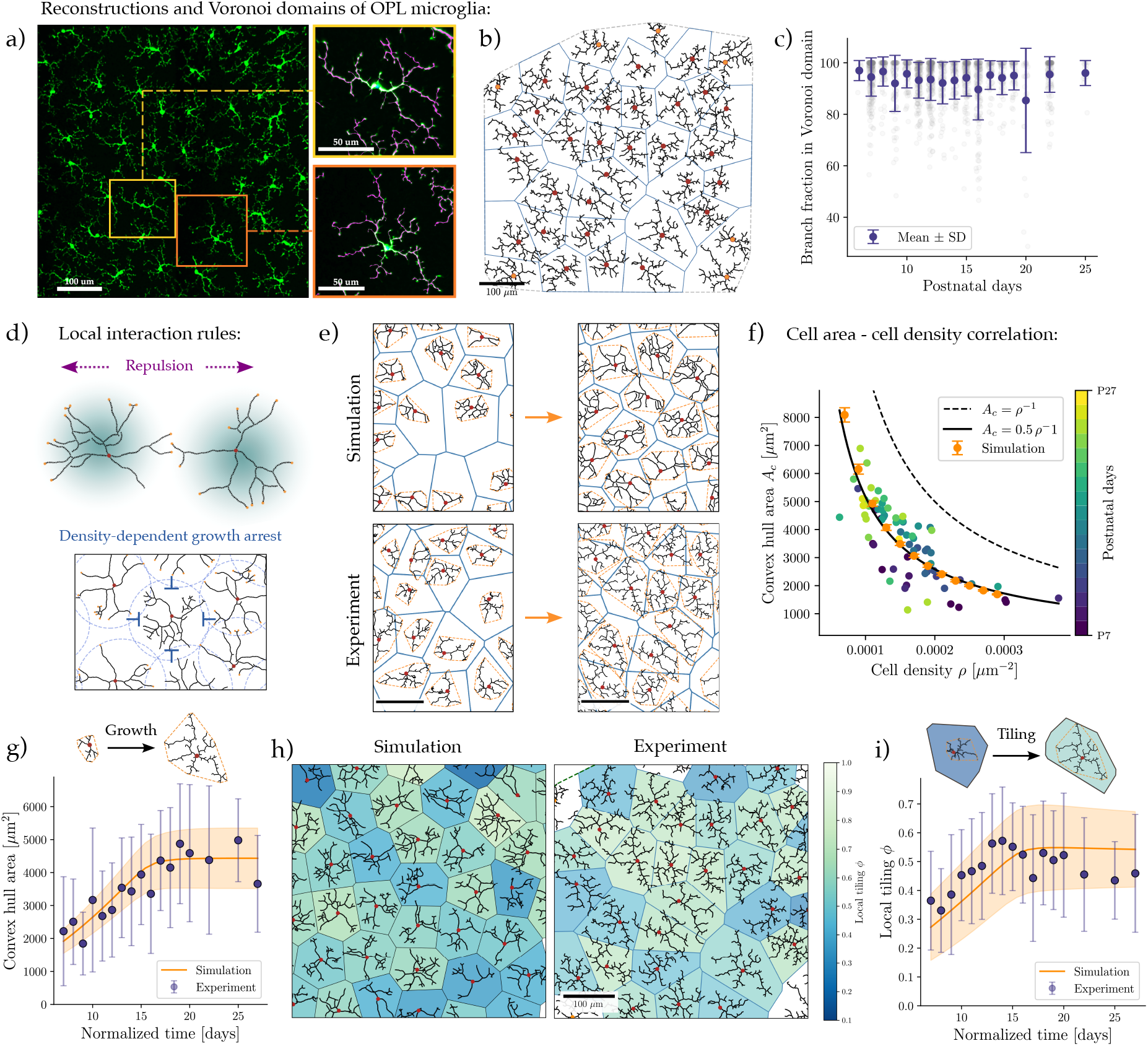
Stochastic branching model with neighbor repulsion and density-dependent growth arrest reproduces microglial patterning in the developing retina. **a)** Confocal images of immunostained microglia (Iba1, green) in the outer plexiform layer (OPL) of adult retina, with reconstructed individual morphologies (zoom). **b)** Voronoi tessellation constructed from soma positions (red) from **a**. Dashed line: Global convex hull of all microglia in the subregion. **c)** Fraction of each cell’s arbor within Voronoi domain (gray dots) across postnatal development. Markers: experimental mean with ± SD. **d)** Schematic of the stochastic branching model. Top: Individual cells grow as branching-and-annihilating random walks, with active tips (orange) that terminate upon contact with other branches. A repulsive interaction field (teal) drives neighboring cells apart. Bottom: Growth arrests when the local area fraction of overlapping cellular territories (dashed circles, defined by maximal branch reach) exceeds the hexagonal close-packing threshold. **e)** Representative simulation (top) and experimental (bottom) snapshots at early (P8, left) and late (P22, right) developmental stages. Dashed orange lines: Convex hull around cellular arbors. Blue outlines: Voronoi territories. Scale bar: 100 *µ*m. **f)** Convex hull area *A*_*c*_ vs. cell density *ρ* across developmental stages (color-coded). Dashed line: scaling predicted for full area coverage (*A*_*c*_ = *ρ*^*−*1^). Solid line: fit to experimental data. Orange dots: simulation predictions. **g)** Developmental growth of individual microglia quantified by convex hull area. Markers: experimental mean with ± SD. Solid line and shaded region: simulation mean ±SD. Simulation time normalized to experimental time points. **h)** Local tiling of microglia (P22) in simulations (left) and experiments (right), color-coded by the metric *ϕ*. **i)** Mean local tiling across development. Markers: experimental mean ±SD. Solid line and shaded region: simulation mean ±SD. *n* = 3–4 animals per age, *N* = 10–60 cells per OPL field.

Reconstructed branching patterns of individual cells exhibited a characteristic combination of stochasticity and regularity: cells varied in branch number, elongation, and maximal reach, yet consistently displayed radially extending, outward-growing architectures (**Fig. 1a**). To account for this interplay, we turned to a mathematical model of stochastic branching that we previously used to describe the morphogenesis of sensory neurons [32]. In this framework, arbors grow as branching and annihilating random walks (BARWs) [43]: active tips elongate and branch stochastically at a fixed rate, terminate upon contact with existing branches, and reorient through self-avoidance or global guidance cues [32] (see **Supplementary Note S1** for model details). To test the applicability of the BARW model, we focused on single-cell statistics from microglia from P7 to P22. First, individual branch lengths were exponentially distributed, as predicted by a stochastic branching process (**Supp. Fig. S1a**). Second, the mean branch length, and thus the branching probability, remained constant across developmental stages, indicating a conserved growth rule (**Supp. Fig. S1a-b**). Third, radial branch orientation distributions matched a peaked Gaussian profile, as predicted by the BARW framework for arbors shaped by a radial guidance cue [32], with a variance that was likewise conserved across development (**Supp. Fig. S1c-d**). Together, these measurements fully parameterized the model, which reproduced the characteristic morphologies of reconstructed microglia when the branching process was terminated at the observed cell sizes (**Supp. Fig. S1e**). However, the single-cell BARW model treats final cell size as an extrinsic parameter, rather than an emergent property. This raises the question of whether, in a tissue with many microglia, cell sizes are collectively regulated through local interactions.

### Local repulsion and density-dependent growth arrest govern collective domain formation

When multiple microglia colonize the same tissue layer, cells may stop growing either when contacting neighbors, sensing local crowding, or independently reaching a target size. To determine which of these mechanisms operates, we first quantified how microglia collectively partition space. In the adult retina, neighboring microglia arbors showed minimal interpenetration (**Fig.1a**), suggesting well-defined domains around each cell. To quantify these domains, we constructed Voronoi tessellations using soma positions as generators, which provides a parameter-free assignment of local territories (**Fig.1b**). For each cell, we then computed the fraction of reconstructed arbor length within its Voronoi polygon. Throughout development, microglial arbors were strongly confined to their assigned territories, with, on average, only 5% of arbor length crossing into neighboring Voronoi domains (**Fig.1c**). Remarkably, individual branches navigate locally with no explicit information about soma-defined Voronoi boundaries, yet rarely cross into neighboring territories.

This tight domain structure places two constraints on collective patterning. (i) First, Voronoi domains must remain sufficiently homogeneous across development. BARW simulations with *N* cells seeded at random positions showed large fluctuations in local density, such that the contact-terminated branching model led to branches in dense regions terminating prematurely while surviving branches found their way into neighboring domains, producing strong arbor crossings and cell size heterogeneity (**Supp. Fig. S2a-c**). In contrast, our data show that the convex hull area *A*_*c*_, which encloses all branches of a cell, displayed a clear peak across development, suggesting that cell sizes were not randomly distributed and had a well-defined mean (**Supp. Fig. S3a**). (ii) Second, the final cell size must be set by a rule beyond tip-branch contact or cell-autonomous growth arrest. Pure contact-mediated termination predicts average cell area scaling as ⟨*A*_*c*_⟩ = *ρ*^−1^, where *ρ* is the overall cell density, and crowding-independent growth arrest predicts no correlation between *A*_*c*_ and *ρ*. Instead, we experimentally observed ⟨*A*_*c*_⟩ ≃ 0.5 *ρ*^−1^ across developmental stages (**Supp. Fig. S3b**), indicating a strong correlation by which cells arrest their growth well before exhausting the available space. Together, these results ruled out both density-independent growth arrest and purely contact-driven branch termination.

To satisfy the first constraint (i), we introduced a repulsive soft-core interaction between neighboring cells (**Fig. 1d**; see **Supplementary Note S1** for details), following recent particle-based simulation frameworks [14, 44]. Somatic translocations of microglia have been reported *in vivo* [45], and analogous repulsive interactions have been observed in other retinal cells such as starburst amacrine cells [30, 46]. Recent work has also identified molecular regulators of microglial tiling whose downstream targets include repulsive receptor pathways [31]. In the absence of detailed information on the molecular coupling between branch contacts, arbor remodeling, and soma translocations, this effective potential provides a parsimonious description of how local interactions redistribute cell positions, analogous to recent cell migration models based on local chemokine secretion [47]. We found that local repulsion homogenized Voronoi domains but did not by itself resolve the branch interpenetration into neighboring territories (**Supp. Fig. S2ac**).

To satisfy the second constraint on the final cell sizes (ii), we introduced a crowding-dependent growth arrest rule. While several biologically plausible mechanisms can be envisioned, such as termination driven by contact-dependent branch retraction or local depletion of growth factors, these require defining additional parameters not constrained by our data. We therefore adopted a minimal geometric criterion: each cell is assigned a spherical “interaction” region defined by the maximal reach of any of its branches, and branching terminates when the local packing fraction of neighboring regions reaches the 2D hexagonal close-packing threshold 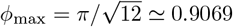 [48] (see **Fig. 1d** and **Supplementary Note S1**). The choice of a spherical region aligned with our observation that sphericity of convex polygons around each cell remained close to 0.8 across development (**Supp. Fig. S3c**).

We found that the coupled stochastic branching model with neighbor repulsion and density-dependent growth arrest reproduced the key statistics of branching patterns and domain segregation at both early and late developmental stages (**Fig. 1e**). Notably, by varying cell density *in silico*, we recovered the experimentally observed scaling *A*_*c*_ ≃ 0.5 *ρ*^−1^ (**Fig. 1f**) as a result of the geometric packing rule without any parameter tuning. After matching time scales between experiment and theory by comparing experimental and simulated convex hull areas, we found that the model captured the temporal evolution of cell sizes accurately across development, which showed a monotonic increase until P19 followed by saturation (**Fig. 1g**).

### Microglia progressively fill their local Voronoi domains until maturation

Having established that individual microglial arbors expand significantly during early development, we next asked whether this growth reflects active space-filling or simply passive kinematic scaling due to isotropic tissue expansion. A purely passive tissue expansion model predicts a proportional increase in both intercellular distances and local Voronoi areas. Instead, we found that up to P16, the mean Voronoi area and global cell density remained approximately constant (**Supp. Fig. S3d**), while individual arbor areas increased more than two-fold (**Fig. 1g**). Combined with minimal invasion into neighboring Voronoi domains (**Fig. 1c**), these distinct geometric constraints indicate that early arbor expansion is driven by the progressive filling of a fixed local territory. Only after P16, Voronoi areas begin to increase alongside a drop in cell density, likely reflecting late-stage tissue expansion or cell death (**Supp. Fig. S3d**). To quantify this local tiling mechanism, we measured for each cell the fraction of its Voronoi domain covered by its arbors. We defined the tiling metric as *ϕ* ≡ *Ā*_*c*_*/A*_*v*_, where *Ā*_*c*_ is the area of the convex hull around the cell’s branches that lies within its Voronoi domain, and *A*_*v*_ is the area of the Voronoi domain. With this definition, *ϕ* = 1 corresponds to complete coverage of the local territory. Applying this measure to simulations and experiments, we found similar spatial patterns of local tiling, with comparable cell-to-cell variability across the tissue (**Fig. 1h**). Next, we quantified the mean tiling metrics over development. Tiling increased monotonically from *ϕ* ≃ 0.3 at P7–P8 to *ϕ* ≃ 0.55 at around P15 both in the model and experiments (**Fig. 1i**), confirming that microglia grow and efficiently tile their local domains during the retina maturation phase. Relaxing either the neighbor repulsion rule or the density-dependent growth arrest *in silico* failed to reproduce this pattern. Cells either oversaturated the region or spread inefficiently into neighboring domains (**Supp. Fig. S2b**,**d**).

### Cell positions exhibit an increasingly pronounced hyperuniform pattern over development

Our model predicts that neighbor repulsion improves local tiling but also redistributes soma positions away from locally crowded regions, which enhances spatial regularity. We therefore asked whether microglial soma positions become increasingly ordered during development. Biologically, achieving such large-scale order prevents voids and dense clusters, ensuring optimal resource allocation and uniform tissue coverage without the constraints of a rigid crystalline lattice. Recently, the notion of hyperuniformity has been developed [34], which provides a quantitative signature of long-range order in biological systems [35, 36, 49]. To test for hyperuniformity, we first measured number fluctuations by counting the number of somata *N* (*R*) inside circular observation windows of radius *R* and computing the variance *σ*^2^(*R*) across *n* = 50 randomly placed windows for each radius (see **Supplementary Note S4**). For an uncorrelated (Poisson) point pattern in two dimensions, 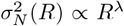 with *λ* = 2. For correlated patterns at large scales, fluctuations grow more slowly with *R*, approaching *λ* = 1 in the crystalline limit. In hyperuniform patterns, long-wavelength fluctuations are anomalously suppressed, corresponding to exponents *λ <* 2 for large *R* (**Fig. 2a**).

**FIG. 2.**
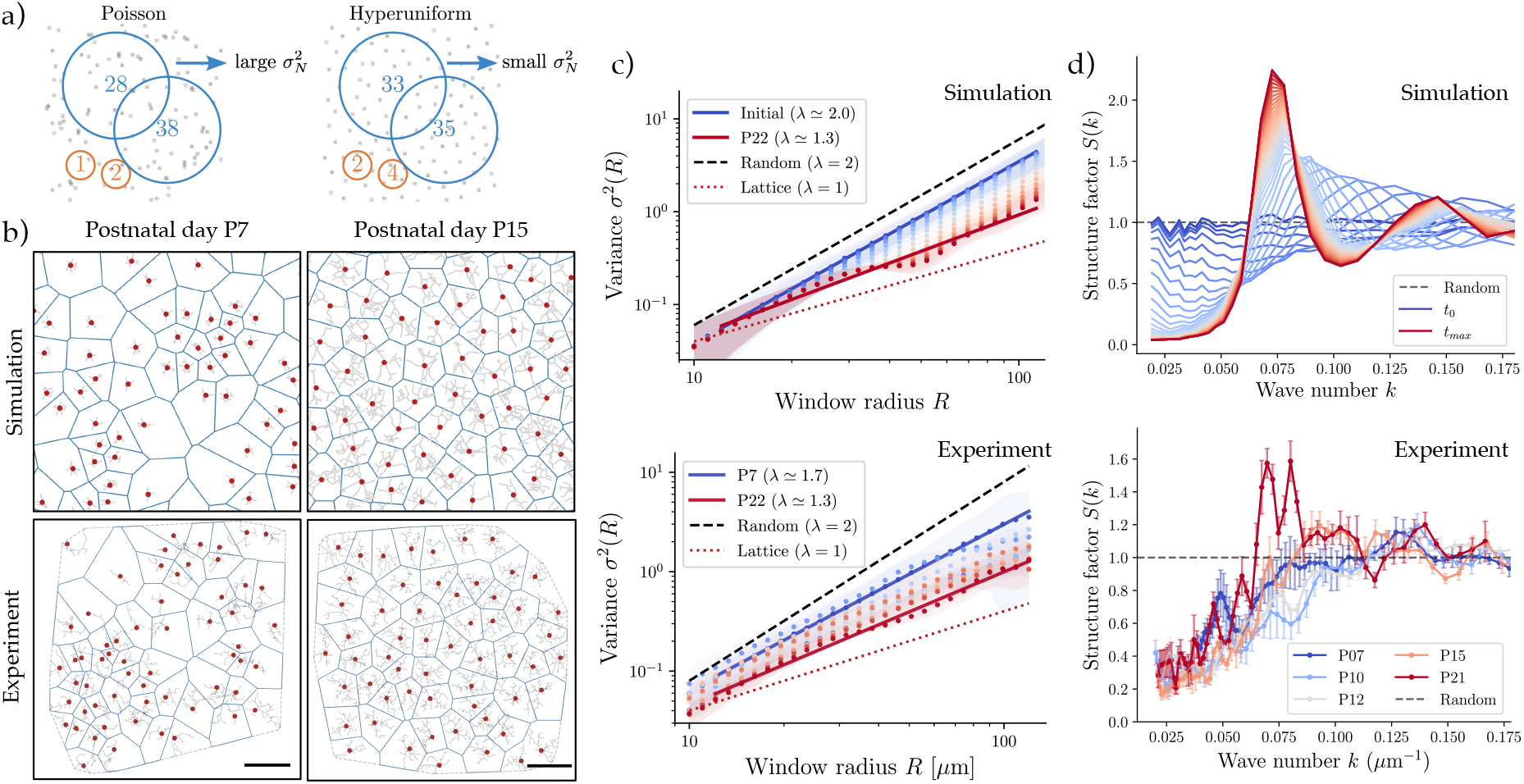
Cell position fluctuations are progressively suppressed during development. **a)** Schematic illustration of hyperuniformity. Circular windows of varying radii (orange, blue) count the number of enclosed particles at different spatial scales for a Poisson (left) and a hyperuniform (right) point pattern. Unlike the Poisson case, the hyperuniform pattern exhibits suppressed variance in particle number for large radii *R*, following a scaling 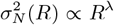 with *λ <* 2. **b)** Representative cell soma coordinates (red dots) from simulations (top row) and experiments (bottom row) at P7 (left) and P15 (right). Blue outlines: Voronoi territories. Scale bar: 100 *µ*m. **c)** Spatial uniformity quantified from the number variance *σ*^2^(*R*) of somata counted within circular observation windows of radius *R*. Top: In simulations, the initial state follows a scaling close to *σ*^2^(*R*) ∝ *R*^2^ (dashed line). At later stages, corresponding to experimental time point P22, the variance scaling shifts toward smaller exponents close to 1 (dotted line). Bottom: Experimental number variance scaling between P7 and P22. Markers: pooled mean values across samples at each *R*. Shaded regions: standard deviation. **d)** Top: Time evolution of the static structure factor *S*(*k*) from simulations. Color code indicates increasing time from initial (blue) to final time point (red). Bottom: Static structure factor *S*(*k*) at different developmental time points (color-coded) from experiments. Dashed line: *S*(*k*) = 1 for a Poisson point process.

Early developmental stages up to P9 exhibited irregular soma arrangements similar to the randomized initial conditions used in our simulations. In contrast, later stages showed a more regular pattern (**Fig. 2b**). Quantitatively, we found that simulations predicted a monotonic reduction of the number-fluctuation exponent from a Poisson scaling (*λ* ≃2) to *λ*≃ 1.3 as repulsive translocations proceed (**Fig. 2c, Supp. Fig. S2e**). Testing this on our experimental data, we found a similar trend of *λ* decreasing from *λ*≃ 1.7 at P7 to *λ*≃ 1.3 by P22 (**Fig. 2c**). The narrow experimental range of *λ*≃ 1.2–1.4 at late stages (**Supp. Fig. S3e**) tightly constrained the effective strength of repulsive interactions in the model, as confirmed by systematically varying this parameter (**Supp. Fig. S2f,g**; see **Supplementary Note S1** for details).

Next, we computed the static structure factor *S*(*k*) of soma positions as a complementary probe in Fourier space, which follows *S*(*k*) → 0 as *k* → 0 for hyperuniform patterns. In simulations, *S*(*k*) evolved from an initial state with *S*(*k*) ≃ 1 at all *k* toward a state with a pronounced peak at intermediate *k* and a clear suppression of *S*(*k*) at the smallest accessible wave numbers (**Fig. 2d**), indicating progressively reduced long-wavelength density fluctuations. Testing this on the experimental data, we found a similar temporal evolution of *S*(*k*), exhibiting a decay for decreasing *k* at later developmental stages (**Fig. 2d**). The structure factor profile at P21 showed a peak at *k* ≃ 0.07 *µ*m^−1^, corresponding to a length scale of approximately 90 *µ*m, comparable to the typical lateral extent of individual microglial territories, matching our theoretical prediction. Together with the number-fluctuation scaling (**Fig. 2c**), these results indicate that microglial soma patterns become increasingly uniform during development, consistent with an approach toward hyperuniform organization over the experimentally accessible length scales.

### Signatures of increasing regularity in cell positions by geometric and topological statistics

To further quantitatively test whether the mechanism of regular patterning arises gradually through local interactions across development, we analyzed the global statistics of soma positions across development. For this, we first focused on the Voronoi domain statistics and the dual Delaunay triangulation of the cell position network, informed by recent studies on hyperuniform Voronoi patterns [50]. We quantified how Delaunay edge lengths, which link two neighboring Voronoi domains, were distributed across development (**Fig. 3a**). In our *in silico* model, we found a transition from a broader distribution with large tails at early times to a narrower edge length distribution over time, which agreed well with both experimental distribution profiles and average values across development (**Fig. 3b**). Because regular or hyperuniform point pattern distributions converge toward a Gaussian shape [50], we next analyzed the normalized Voronoi area distribution by the mean Voronoi domain area across development. Both simulations and experiments showed large tails in the initial phases and gradually approached a near-Gaussian shape at later developmental times (**Fig. 3c**). Moreover, we found that the regularity index, defined by the ratio of mean to standard deviation of the Voronoi domain statistics [51], increased progressively over development (**Supp. Fig. S3f**).

**FIG. 3.**
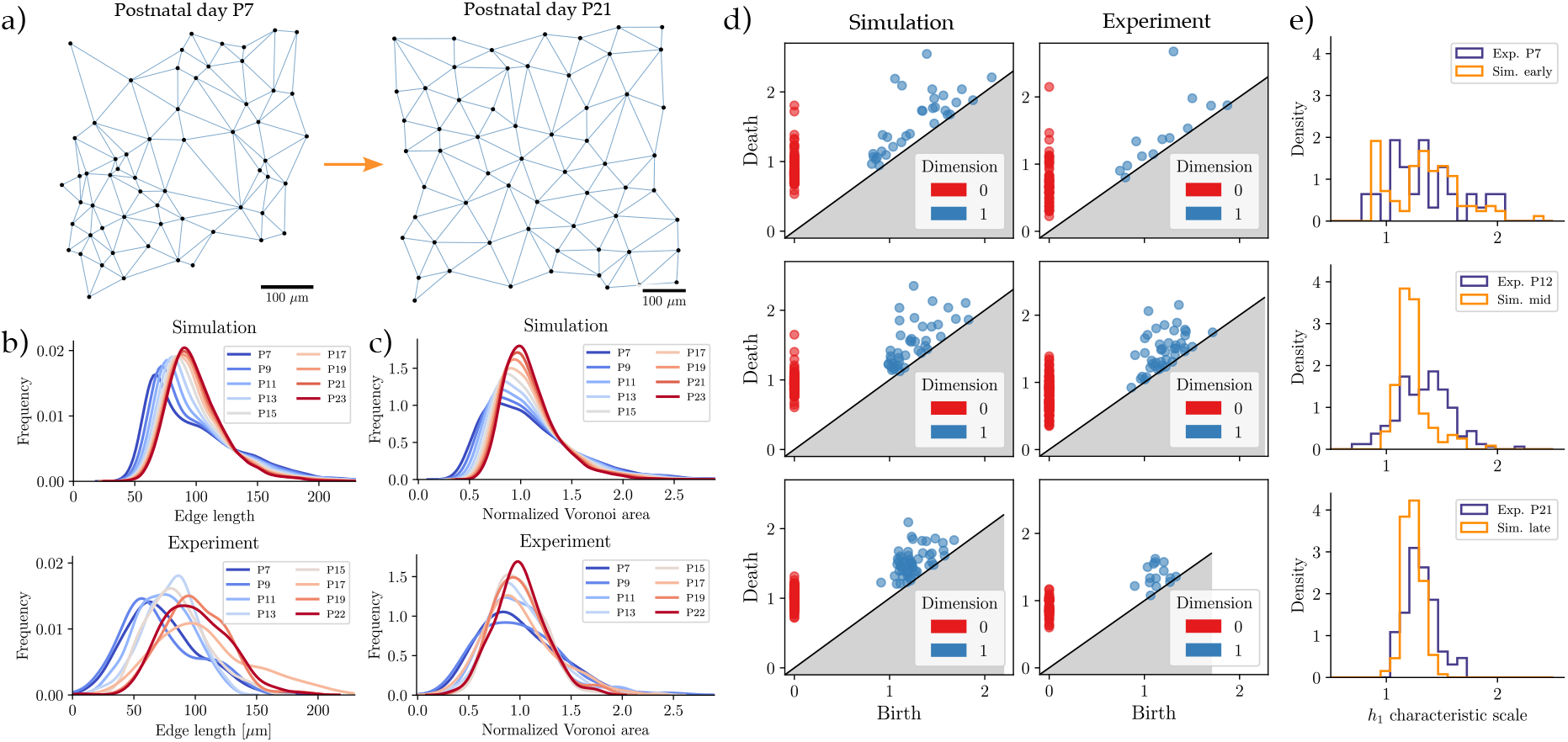
Geometric and topological statistics of soma positions reveal increasing spatial regularity during development. **a)** Representative soma positions (dots) at early (P7) and late (P21) developmental stages, with the Delaunay triangulation representing links between neighboring cells (solid lines). **b)** Distributions of Delaunay edge lengths corresponding to neighbor-neighbor distances in simulations (top) and experiments (bottom) at different time points (color-coded). **c)** Voronoi cell areas normalized by the mean Voronoi area in simulations (top) and experiments (bottom) at different time points (colorcoded). **d)** Persistent homology analysis of soma positions across developmental stages. Representative persistence diagrams with birth and death of connected components *h*_0_ (dimension 0, red) and holes *h*_1_ (dimension 1, blue) as the union of growing domains around soma positions is expanded. Simulation (left) and experimental (middle) persistence diagrams over P7, P12, and P21 (top to bottom). **e)** Pooled distributions characteristic scale of holes *h*_1_ for P7, P12, and P21 (top to bottom).

As regular cell positions can also arise in ordered non-hyperuniform point patterns like in a random sequential addition process [48, 52], we further tested hyperuniformity using alternative metrics on cell positions. Topological data analysis is an emerging method to capture key properties in point data, including applications in synthetically generated hyperuniform patterns [53]. Specifically, persistent homology captures the structure of point patterns across spatial scales by growing disks of radius *r* centered at each point and tracking how topological features appear and disappear as *r* increases [54]. For example, when two disks first touch, they form an edge merging two connected components, marking the death of an *h*_0_ feature. When three or more disks connect to surround a gap, a hole or loop *h*_1_ is “born”. As *r* grows further, these features eventually merge or give rise to higher-dimensional features, i.e. they “die” (see **Supp. Fig. S4**a for an illustration). The birth and death radii of each feature encode characteristic scales in the pattern (**Supplementary Note S2**).

To examine whether in our system similar signatures of hyperuniformity arise, we first applied persistent homology to the temporal evolution of simulated cell positions. To enable comparison across simulations and experimental samples with different boundary conditions and cell numbers, we normalized all position coordinates by the mean inter-soma spacing (see **Supplementary Note S4**). Constructing persistence diagrams at different time points, we found that both connected components (*h*_0_) and holes (*h*_1_) attained increasingly narrow distributions of birth and death radii over time (**Fig. 3d**). Intuitively, this narrowing reflects reduced density fluctuations, which is a key signature for hyperuniformity as observed previously [53]. Turning to experimental data at representative developmental stages, we observed the same trend in the persistence diagrams, indicating that cell positions become increasingly consistent with hyperuniform organization over development, in agreement with the fluctuation analysis (**Fig. 3d, Supp. Fig. S4b** for the full experimental data). To quantify this further, we analyzed the characteristic scale of *h*_1_ features, defined as (birth + death)/2, the midpoint of the filtration interval during which a hole exists. For a regular pattern, *h*_1_ features cluster around a characteristic spacing scale; for a random pattern, this distribution is broader. Pooling across samples, we found that characteristic scale distributions narrowed markedly over development (**Fig. 3e**), further supporting the approach to regular patterning.

### Perturbed microglial morphogenesis disrupts local tiling and large-scale regularity

Our theoretical model predicts that local tiling and large-scale regularity emerge from the interplay of self-organized branching and neighbor repulsion. To test this, we examined microglia in the *Pde6b*^*rd10*^ mouse model [55–57], in which genetically-induced progressive photoreceptor degeneration beginning at P16 triggers microglial proliferation and reduces branch complexity in the OPL [58, 59]. This perturbation alters both cell density and individual morphodynamics, providing an opportunity to assess their respective contributions to spatial order.

At early stages (P10), microglial morphologies and spatial distributions were similar between *rd10* and WT, as expected (**Fig. 4a**). When we looked at microglial patterns after P20, however, the two conditions started to diverge (**Fig. 4a**). WT microglia expanded to fill their local territories, whereas *rd10* cell sizes, quantified by convex hull areas, did not exceed P10 levels until P50 (**Fig. 4b, Supp. Fig. S5a**), indicating arrested arbor expansion. Meanwhile, *rd10* cell density increased from P10 onward, reaching approximately threefold higher values than WT by P50 (**Supp. Fig. S5b**) with correspondingly smaller Voronoi domain areas (**Fig. 4c, Supp. Fig. S5c**).

**FIG. 4.**
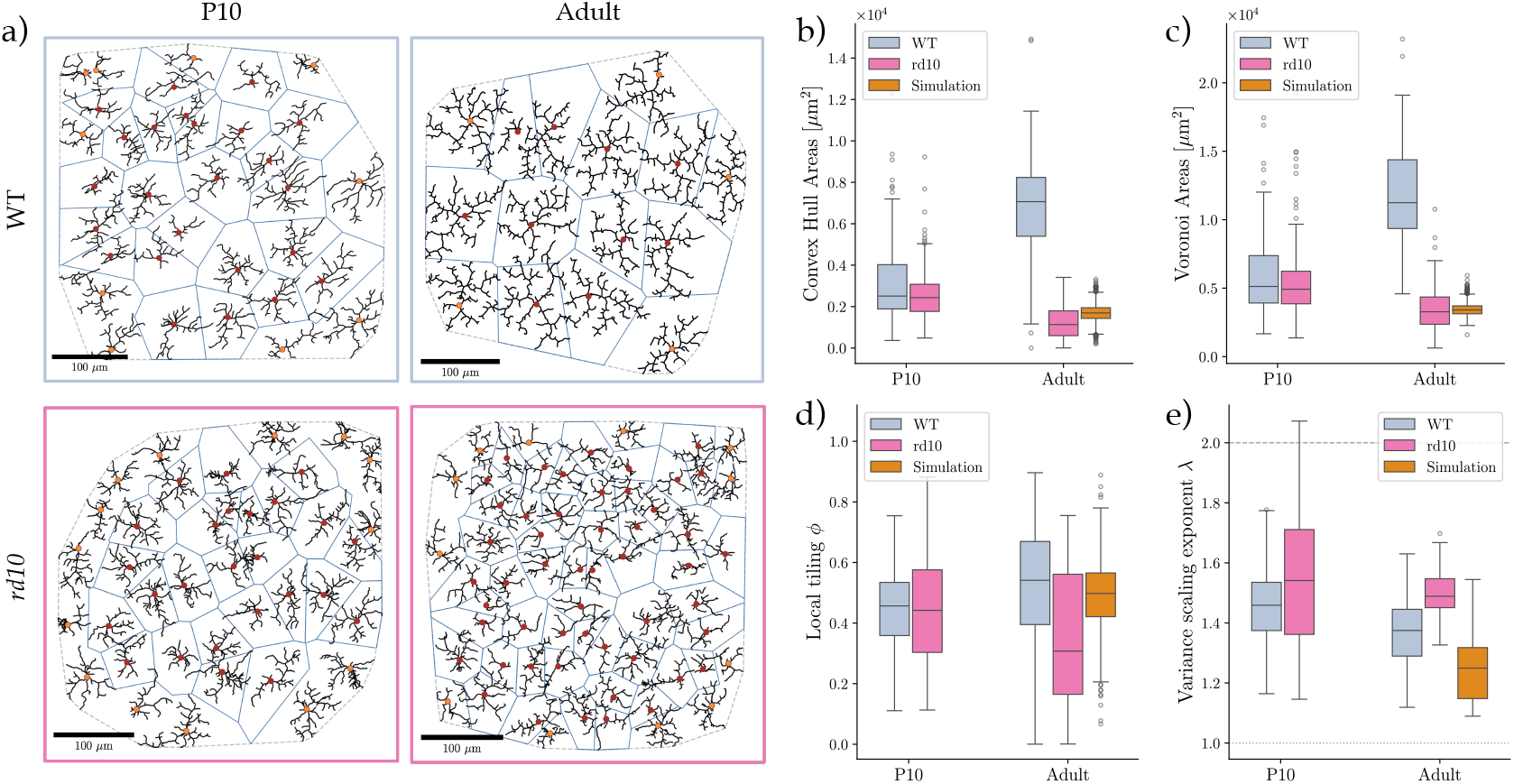
Perturbing microglial morphogenesis impairs developmental tiling and large-scale spatial regularization. **a)** Representative reconstructed microglia morphologies from wild-type (WT; top) and *rd10* (bottom) mice at postnatal day 10 (left) and P22 (right). Red dots: soma with branches (black). Blue: Voronoi domains. **b)** Box plot of cell convex-hull areas from WT (gray), *rd10* (pink), and simulation (orange) data at selected developmental times. **c)** Box plot of Voronoi-area distributions. **d)** Box plot of local tiling efficiency *ϕ*. **e)** Variance scaling exponent *λ* for soma positions. Box plots show median (center line), interquartile range (box), whiskers (1.5× IQR), and outliers (individual points).

To disentangle the effects of increased density from impaired morphodynamics, we used our *in silico* model with cell density matched to the *rd10* adult level, but with the WT branching rules intact. In these simulations, local tiling *ϕ* increased and number fluctuations decreased during growth, following the WT trend (**Supp. Fig. 4d,e**). Increased density and smaller cell sizes alone were therefore not sufficient to disrupt spatial order. In contrast, *rd10* microglia decreased local tiling after P20, reaching *ϕ*≃ 0.3 at P50 compared to *ϕ*≃ 0.55 in the WT adult (**Fig. 4d, Supp. Fig. S5f**). This decrease in tiling occurred even though *rd10* Voronoi domains were approximately three times smaller than WT domains, meaning that less arbor expansion is needed to achieve comparable tiling.

To assess large-scale order, we then computed variance scaling exponents *λ* from soma positions. The *rd10* developmental trajectory was noisier than the WT case, with intermediate stages (P20, P44) showing transiently low *λ* values (**Supp. Fig. S5g**). Nonetheless, the overall trend showed increasing *λ*, reaching *λ* ≃ 1.5 at P50, compared to *λ*≃ 1.3 in the WT adult (**Fig. 4e**). In the density-matched *in silico* model, both tiling (*ϕ* ≃0.5) and fluctuation suppression (*λ* ≃1.25) remained close to WT levels (**Fig. 4d,e**), reinforcing that impaired spatial regularity in *rd10* arises from disrupted morphodynamic interactions rather than from the altered density alone.

### A minimal model of growing repulsive cells links efficient tiling to hyperuniformity

Finally, to isolate the minimal ingredients that jointly promote (i) high territorial coverage and (ii) suppressed long-wavelength density fluctuations, we introduced a coarse-grained Langevin model. Each cell is represented by a growing soft disk of radius *R*_*i*_(*t*) whose center **r**_*i*_(*t*) moves under short-range repulsive interactions (**Fig. 5a**), following an approach based on Hertzian elastic contact theory previously used to describe growing bacterial colonies [60]. Growth is self-limited by local crowding: overlapping contacts inhibit further radial expansion, capturing a generic contact-dependent growth arrest without requiring specification of microscopic details.

**FIG. 5.**
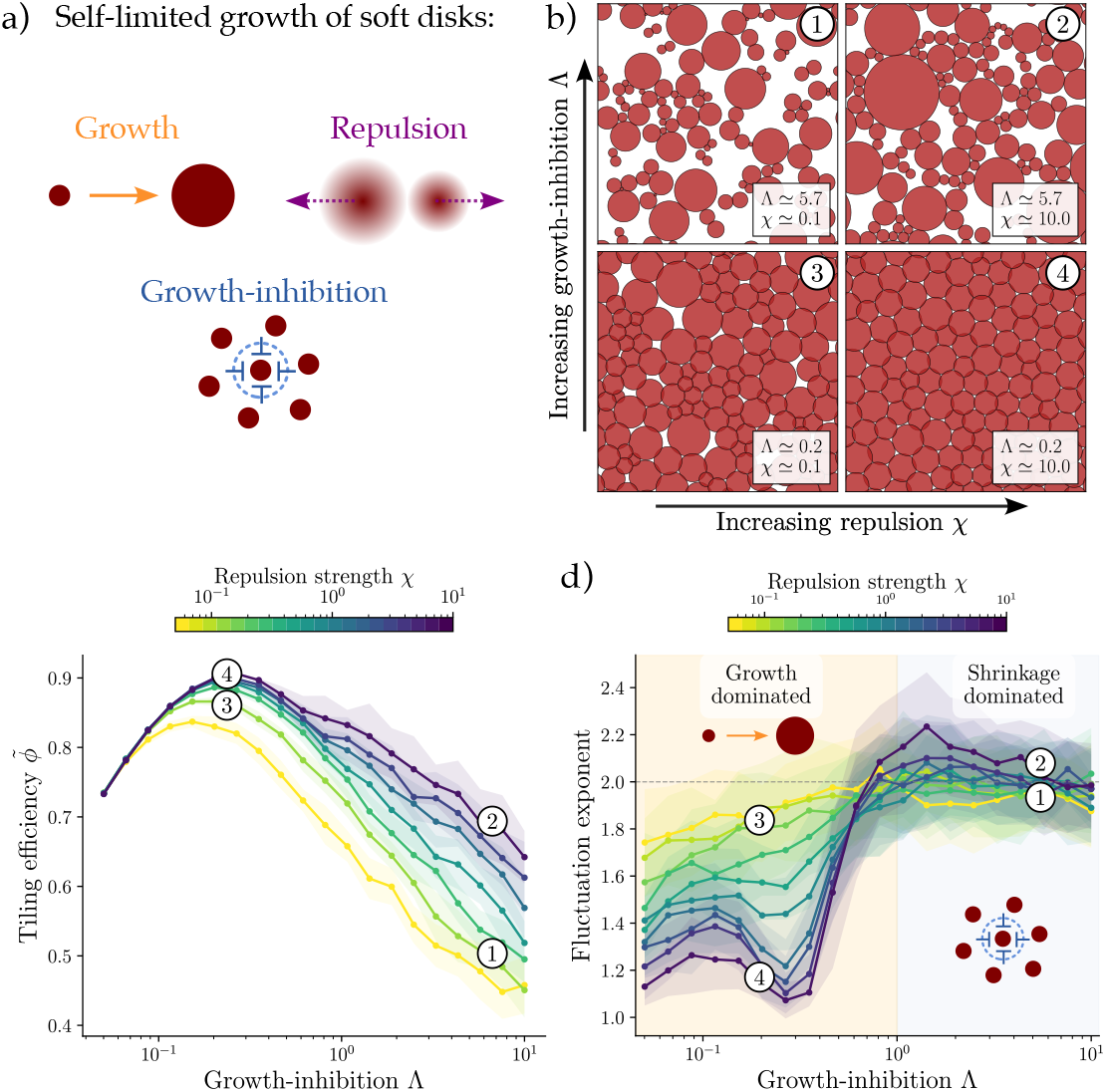
A coarse-grained Langevin model of growing soft repulsive disks captures the mechanism of hyperuniform tiling. **a)** Schematic of the model. Soft disks (red) grow at a uniform rate (orange) and repel each other on contact (purple). Growth is self-limited by shrinkage rate proportional to disk–disk overlaps from neighbors. **b)** Representative patterns across the (*χ*, Λ) parameter plane, where *χ* controls repulsive mobility and Λ controls growth inhibition. For large Λ (top row), disks arrest growth early and coverage remains inefficient. For smaller Λ (bottom row), coverage improves and disk sizes become more homogeneous; increasing *χ* further regularizes the pattern. **c)** Tiling efficiency 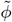 over growth inhibition for different values of the repulsion strength *χ* (color-coded). Tiling efficiency shows a maximum for Λ*≃* 0.2*−*0.4 (Regions 3 and 4). **d)** Phase map of the number-fluctuation exponent *λ* as a function of *χ* and Λ. When shrinkage dominates over growth (Λ *>* 1), *λ* remains close to 2 for all *χ*, consistent with Poisson-like fluctuations (Regions 1 and 2). In the growth-dominated region (Λ *<* 1), sufficiently large repulsion drives *λ* toward 1 (Region 4), indicating strong suppression of long-wavelength fluctuations. When repulsion is weak, exponents remain close to *λ ≃* 2 even at low Λ (Region 3).

After rescaling lengths by the initial radius *R*_0_ and time by the growth time *T*_0_ = *R*_0_*/G* with an intrinsic growth speed *G*, the model takes the dimensionless form

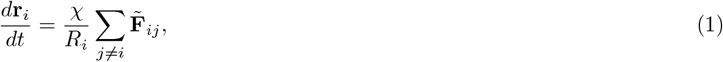

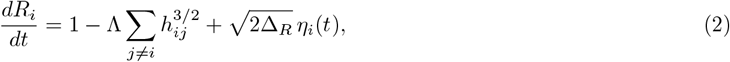

where *h*_*ij*_ is the overlap between cells *i* and *j*, 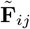 is a dimensionless soft-core repulsive force (see **Supplementary Note S3**), Δ_*R*_ sets the radial fluctuation strength, and *η*_*i*_(*t*) are independent Gaussian white noises. The dynamics is then controlled by the dimensionless parameters,

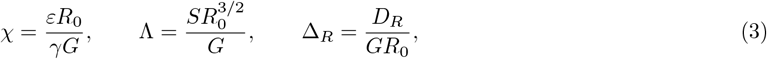

together with the initial density *ρ*_0_. Here, *χ* quantifies mechanical repulsion strength relative to growth with drag coefficient *γ* and elastic modulus *ε*, while Λ is the ratio of contact-induced shrinkage with rate *S* to growth speed *G*: Λ *>* 1 implies that contacts cause net shrinkage, whereas Λ *<* 1 allows net growth even under crowding.

Numerical simulations revealed distinct regimes in the (*χ*, Λ) plane (**Fig. 5b**). For strong inhibition (top row), growth arrests early and the pattern shows insufficient coverage. Under weak repulsion, cells become kinetically trapped in clusters; under strong repulsion, cells in crowded regions shrink while those in sparser regions continue to grow, leading to size heterogeneity and persistent gaps. Reducing Λ (bottom row) promotes more uniform cell sizes and improved surface coverage. However, if inhibition is too weak, disks continue to grow in crowded regions and significant overlaps accumulate, increasing redundancy. Thus, efficient tiling requires a balance: repulsion redistributes cells to suppress clustering, while growth inhibition prevents excessive overlap.

We quantified this trade-off with a tiling efficiency metric 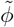, defined as the covered fraction of the surface multiplied by an overlap penalty (see **Supplementary Note S3**). Intuitively, 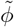 is close to unity only when disks cover nearly the entire surface with minimal mutual overlap; it decreases if coverage is incomplete or if large overlaps create redundancy. Mapping 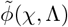 revealed a clear optimum at intermediate Λ, where gaps are suppressed via growth while overlaps remain small (**Fig. 5c**). We found that increasing *χ* generally improves tiling by redistributing disks away from crowded regions, in particular when shrinkage is dominated over growth (Λ *>* 1).

To assess hyperuniformity, we then computed the number-fluctuation scaling 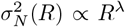 for disk centers. We observed a key transition controlled by the growth-inhibition parameter Λ (**Fig. 5d**). When shrinkage dominates (Λ *>* 1), the fluctuation scaling exponent remains near the Poisson value (*λ* ≃ 2) regardless of repulsion strength *χ*. In contrast, when growth dominates (Λ *<* 1), the exponent drops sharply to *λ* ≲ 1.2 for sufficiently strong repulsion, consistent with a transition toward hyperuniform organization. Notably, the parameter region where *λ* reaches a minimum coincides with the maximum of 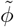, indicating that the same mechanism—strong repulsion to homogenize positions coupled with self-limited growth to prevent redundant overlap—optimizes both tiling efficiency and large-scale uniformity.

## DISCUSSION

Here, we combined a biophysical model for multicellular branching morphogenesis with experiments on microglial patterning in the developing retina. We found that coupling local neighbor repulsion to self-limited growth is sufficient to produce efficient tiling and progressive suppression of large-scale number fluctuations, indicating the emergence of hyperuniformity. Using Voronoi-based measures (**Fig. 1**), fluctuation analysis (**Fig. 2**), and persistent homology (**Fig. 3**), we quantified this transition experimentally during retinal maturation. We furthermore showed that perturbing microglial morphodynamics reduces tiling and long-range order (**Fig. 4**), supporting the idea that sufficient single-cell growth is required to reach the hyperuniform tiling regime.

Our coarse-grained model reveals a transition from Poisson-distributed arrangements to hyperuniform order controlled by the balance between growth and neighbor-induced shrinkage (**Fig. 5**). When shrinkage dominates growth, large-scale correlations follow Poisson statistics; when growth dominates with sufficient cell-cell repulsion, the pattern approaches strong hyperuniformity. This growth-repulsion feedback belongs to a broader class of systems in which size dynamics drive collective organization, including pulsating active systems [61, 62], catalytically active colloids [63], and repulsive microtubule asters [64]. By contrast, *hyperdisorder* has recently been reported in squid chromatophores, where cell proliferation during tissue expansion enhances density fluctuations [65]. In our system, density fluctuations are progressively suppressed under conserved cell number through local repulsive interactions. In a related biological context, astrocyte tiling has been modeled by placing somata sequentially while rejecting overlapping configurations [66]. Although this approach captures territorial exclusion, it is based on a random sequential addition process and therefore cannot produce long-range order [34]. Our study provides a growth-driven route for the emergence of hyperuniformity, and complements recent theoretical work on this, including random reorganization models [67], active particle systems [68, 69], and vertex models of cell monolayers [49].

Biologically, the ordered architecture of the outer retina raises the question of whether microglial spatial regularity passively reflects the underlying photoreceptor mosaic. However, several observations argue against this interpretation. The photoreceptor mosaic is largely established by P5–P7, with densities approaching adult levels by P10–P14 [70, 71]. Yet we observe continued refinement of spatial patterning beyond this window. Moreover, photoreceptor degeneration-induced remodeling in *rd10* mice alters microglial arbor growth and disrupts late-stage fluctuation suppression, suggesting that large-scale spatial regularity is governed by microglial morphodynamics rather than passively inherited from the photoreceptor lattice. This is consistent with the functional role of microglia in locally surveilling the neuronal environment.

Our framework raises several questions for future investigation. First, we have assumed a conserved cell number in our model, yet wild-type microglial density decreases after P16, possibly through apoptosis or tissue expansion [72]. If cell elimination is coupled to local coverage, for instance by selectively removing cells that fail to reach a threshold size, this would introduce an additional optimization linking cell fate to territorial efficiency. Second, microglia interact with neurons [73] and vasculature [74] during development. Whether such heterotypic interactions contribute to the early suppression of density fluctuations, or whether spatial order propagates across cell types through bidirectional coupling, remains open. Layer-specific patterning between the OPL and IPL, where microglia encounter distinct synaptic environments, offers a natural setting to investigate this. Third, long-term *in vivo* imaging of retinal microglia would be partially possible with adaptive optics scanning laser opthalmoscopy [75–77], yet not feasible in juvenile mice prior to eye opening. Therefore, the kinetics of branch retraction, polarity changes, and soma repositioning [7, 23, 24] are not directly observable in this regime, and represent an open direction for future work. In summary, our work suggests that local growth-repulsion feedback may constitute a generic route to ordered tissue coverage wherever growing cells must partition space without centralized coordination.

## MATERIALS AND METHODS

### Animals

Mice were purchased from The Jackson Laboratories: For the developmental dataset, B6.129P-*Cx3cr1* ^tm1Litt/J^ (Cat#005582, named here *Cx3cr1* ^GFP/-^) homozygous mice were crossed with C57BL/6J (Cat#000664). For retinal degeneration, B6.CXB1-*Pde6b*^rd10/J^(Cat#004297, named here *rd10*) founder animals, which have a missense point mutation in exon 13 of the *β*-subunit of the rod cGMP phosphodiesterase gene (*Pde6b*) [78], were backcrossed to the C57BL/6J background for at least 10 generations and then used homozygous. Mice were housed in groups of three to five in the ISTA Preclinical Facility with a 12-hour light-dark cycle. Food and water were provided *ad libitum*. All experiments were performed during the light cycle. All animal procedures are approved by the Bundesministerium für Wissenschaft, Forschung und Wirtschaft (bmwfw) Tierversuchsgesetz 2012, BGBI. I Nr. 114/2012, idF BGBI. I Nr. 31/2018 under the numbers 2021-0.607.460, 2023-0.250.844, 2025-0.781.719, 2020-0.272.234, and BMWFW66.018/0005-WF/V/3b/2016.

### Retina tissue collection

Following cervical dislocation and decapitation, eyes were enucleated with curved forceps. Retinas were rapidly dissected in 1× phosphate-buffered saline (PBS) and transferred to 4% (w/v) paraformaldehyde in PBS (SigmaAldrich, P6148-1KG) for 30 min fixation. After 3× wash in 1× PBS, retinas were placed in 30% (w/v) sucrose (Sigma-Aldrich, 84097-1KG)/1× PBS overnight at 4°C. After three freeze-thaw cycles on dry ice, retinas were washed three times with 1× PBS. Retinas were incubated in blocking solution containing 1% (w/v) bovine serum albumin (Sigma, Cat#A9418), 5% (v/v) Triton X-100 (Sigma, Cat#T8787), 0.5% (w/v) sodium azide (VWR, Cat#786-299), and 10% (v/v) serum (either goat, Millipore, Cat#S26, or donkey, Millipore, Cat#S30) for 1 hour at room temperature on a shaker. Afterwards, the samples were immunostained with primary antibodies diluted in antibody solution containing 1% (w/v) bovine serum albumin, 5% (v/v) triton X-100, 0.5% (v/v) sodium azide, 3% (v/v) goat or donkey serum, and incubated for 48 hours on a shaker at room temperature. The following primary antibodies were used: goat *α*-Iba1 (Abcam, ab5076, Lot FR3288145-1, 1:250) and rabbit anti-Iba1 (GeneTex, Cat#GTX100042, Lot 41556, 1:750). The slices were washed 3 times with 1× PBS and incubated in the secondary antibodies diluted in antibody solution on a shaker, protected from light for 2 hours at room temperature. The secondary antibodies raised in goat or donkey were purchased from Thermo Fisher Scientific (Alexa Fluor 488, Alexa Fluor 568, Alexa Fluor 647, 1:2000). The retinas were washed 3× with 1× PBS. The nuclei were labeled with Hoechst 33342 (Thermo Fisher Scientific, Cat#H3570, 1:5000) diluted in 1× PBS for 15 minutes at room temperature. After washing, the retinas were mounted on microscope glass slides (Assistant, Cat#42406020) with coverslips (Menzel-Glaser #0) using an antifade solution [10% (v/v) mowiol (Sigma, Cat#81381), 26% (v/v) glycerol (Sigma, Cat#G7757), 0.2M tris buffer pH 8, 2.5% (w/v) Dabco (Sigma, Cat#D27802)].

### Confocal microscopy

Images were acquired with a Zeiss LSM880 upright Airy scan or with a Zeiss LSM700 upright using a PlanApochromat 40× oil immersion objective N.A. 1.4. 2 × 2 z-stack tail-images were acquired with a resolution of 1024 × 1024 pixels. For experiment in Figures 2d and 3d-e, retinal whole mounts were imaged using a Nikon confocal microscope (Dual Prime BSI-CSU) equipped with a spinning disk confocal scanner (25 µm pinhole) and a 20× objective lens. Image acquisition was performed with a 10% tile overlap and an exposure time of 70 ms per channel.

### Image processing

Confocal tile images were stitched using the software Imaris Stitcher 9.3.1.v. Then, the confocal images were loaded in Fiji 1.52e (http://imagej.net/Fiji). To remove the background, the rolling ball radius was set to 35 pixels, and images were filtered using a median 3D filter with x, y, z radii set at 3. Image stacks were exported as .tif files, converted to .ims files using the Imaris converter, and imported into Imaris 8.4.2.v. (Bitplane Imaris). For Nikon images, .nd2 format were converted as N5/XML using the python function create bdv n5 multi tile from the python package bdv tools.bdv creation developed by the ISTA Imaging and Optics Facility (IOF). Following image conversion, individual image tiles were stitched using the BigStitcher plugin in FIJI (ImageJ). After tile stitching, image fusion was performed, and the channel corresponding to microglia labeling (Iba1, 488nm) was isolated. The resulting images were saved as TIFF files.

### Reconstruction of 3D-traced microglia

After filtering and background subtraction, images were imported in Imaris 9.2.v (Bitplane Imaris). Microglial processes were 3D-traced with the filament-tracing plugin. Since the filament-tracing plugin provides a semi-automated reconstruction, this eliminates the need for a user-blind approach for selecting representative microglia. New starting points were detected when the largest diameter was set to 12 µm and with seeding points of 1 µm. Disconnected segments were removed with a filtering smoothness of 0.6 µm. After the tracing, we manually removed cells that were sitting at the border of the image and were only partially traced so that these cells would not be analyzed. The generated skeleton images were converted from .ims format (Imaris) to .swc format 157 by first obtaining the 3D positions (x,y,z) and the diameter of each traced microglial process using the ImarisReader toolbox for MATLAB (https://github.com/PeterBeemiller/ImarisReader) and then exporting for format standardization using the NL Morphology Converter (http://neuroland.org). Artifacts from the 3D-reconstructions automatically failed to be converted into .swc format.

### Model details and data analysis

Full details of the simulation framework, including parameter values, model sensitivity analysis, and the coarsegrained Langevin model, are provided in the **Supplementary Note**. The persistent homology analysis pipeline and the reconstruction and statistical analysis of experimental data are also described there.

## CODE AND DATA AVAILABILITY

Custom made scripts to reproduce the findings of this study will be made publicly available in the following Github link: https://github.com/mehmetcanucar/. Source data obtained from the simulations and experiments that support the findings of this study will be provided in the Zenodo repository.

## ACKNOWLEDGMENTS

We thank the scientific service units at ISTA, especially the imaging optic facility (IOF), the preclinical facility (PCF), specifically Michael Schunn and Sonja Haslinger; the Siegert team members for constant feedback on the project, especially Alessandro Venturino, Mohammadhossein Faramarzi, Gloria Colombo, Rouven Schulz, Thomas Negello; Edouard Hannezo for discussions and feedback on the manuscript.

## AUTHOR CONTRIBUTIONS

Conceptualization: SS, MCU; Methodology: SS, LK, MCU; Theoretical model: MCU; Formal Analysis: MCU, LK (TDA); Writing – Original Draft: SS, MCU (lead); Visualization: SS, LK, MCU; Funding acquisition: SS. All authors reviewed and approved the final manuscript.

## Supplementary Note – Self-organized tiling generates tissue-scale hyperuniformity during development

### Supplementary Section S1: Simulation details

#### S1.1. BARW model for individual cells

We model the growth of a single microglial cell as a two-dimensional branching and annihilating random walk (BARW) [43], based on our study of neuronal morphogenesis as a stochastic branching process under local and global guidance signals [32]. Here we briefly summarize the main rules underlying the BARW framework. The growing cell is represented as a set of active tips that perform a persistent random walk, and leave behind inactive branch segments that remain immobile. An active tip gives rise to two new active tips with a fixed probability of branching *p*_*b*_, which sets the average branch length to be ∼ 1*/p*_*b*_. Tips grow on a 2D plane in discrete time steps of fixed length 𝓁 = 1 (in simulation units). The simulation runs for a fixed number of steps *T*_max_, which sets the total spatial extent of the branched cell and serves as a proxy for cell size; no boundary conditions are imposed. Because we do not have experimental access to detailed tip-branch interactions, here we focus on the minimal set of rules to reproduce microglial morphologies and do not consider self-avoidance of branches upon contact, unlike the full phase diagrams explored in [32].

##### Elongation

At each time step, an active tip elongates with probability 1 − *p*_*b*_. The polarity angle of the tip is updated as *φ* → *φ* + *δφ*, where *δφ* is drawn uniformly from [0, *φ*_*e*_]. *φ*_*e*_ = *π/*10 is the maximum angular deviation per step, controlling the persistence of the walk.

##### Branching

With probability *p*_*b*_, a tip bifurcates into two new progeny tips and becomes inactive itself. Each new tip is placed one step 𝓁 from the bifurcation point at angles *φ* ± *φ*_*i*_, where each *φ*_*i*_ is drawn independently and uniformly from [*π/*5, *π/*2] to allow for a minimal separation between the two siblings upon branching.

##### Annihilation

To implement contact-mediated tip termination events, we use an annihilation rule for active tips. After each elongation or branching event, an active tip becomes inactive if it falls within a distance *r*_ann_ of any existing node in the network. The tip’s own parent branch and its sibling branch (specifically, the 2*r*_ann_ most recent nodes of the parent and the first *r*_ann_ nodes of the sibling) are excluded from this check to prevent premature annihilation near the originating bifurcation point.

##### Radial guidance

In addition to the unbiased BARW framework, branches can be influenced by guidance cues acting on the polarity of growing tips. Indeed, sensory neuron morphogenesis has been found to follow a radial guidance cue [32], which directs branches outward from the soma position. As we discuss in Section S4 S4.1 below, we also found a non-zero radial bias controlling the morphologies of individual microglia in this work. This indicates the presence of a radial cue orienting the branches outward from the soma position.

Here we briefly describe the rules underlying this guided BARW framework, following [32]. Each active tip is subject to a radial guidance cue directed outward from the soma. Rather than applying a post-hoc positional displacement, guidance is incorporated as a bias term directly into the stepping probabilities of the underlying random walk [32]. At each step, we define *ψ* to denote the angular difference between the tip’s current polarity angle *φ* and the local radial outward direction 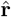. The two possible outcomes during an elongation event, with their corresponding probabilities are:

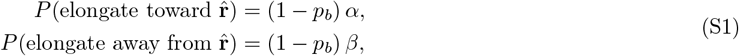

Where

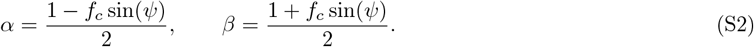

The parameter *f*_*c*_ sets the strength of the radial guidance cue, with *f*_*c*_ = 0 recovering unbiased growth. Because a bifurcation generates two new tips with angular jumps into opposite sides of the parent’s polarity direction, we do not implement the bias term for branching events, i.e. *P* (branch, forward-biased)= *P* (branch, backward-biased)= *p*_*b*_. Since *α* + *β* = 1, total probability is conserved at each step.

##### Parameter estimation from experimental morphologies

The branching probability *p*_*b*_ and guidance strength *f*_*c*_ are estimated directly from SWC-format morphological reconstructions of experimental microglia, using branch segment statistics and radial angle distributions as described in Section S4 S4.1. The estimated values used in all simulations are listed in Table S1.

#### S1.2. Details of the *N* -cell branching model

Here we briefly describe the extension of the BARW framework to describe the collective patterning of an arbitrary number *N* of growing and interacting branched cells.

##### Initialization

To simulate the growth of branched cells on a two-dimensional surface, we place *N* cells with soma positions drawn uniformly at random within a square domain of side length *L* and impose periodic boundary conditions. Each cell is seeded with a small number of primary branches (drawn randomly from 3 or 4), generated by a short persistent random walk from the soma. From this initialization, each cell evolves independently according to the single-cell BARW rules described above, with radial guidance directed outward from its own soma.

##### Cell-cell repulsion

A central feature of our experimental observations on microglial development is that cells attain increasingly regular distributions of their soma positions. To model this, we assume that cells redistribute their soma positions in response to neighboring cells. Biologically, such an interaction could be driven by, e.g., repulsive diffusible cues secreted by each cell, or sequential contact-mediated branch retraction and regrowth dynamics. To capture this at the coarse-grained level, we introduce a soft repulsive interaction between neighboring cells that is proportional to cell size. This effective description provides a simple rule to model repulsive intercellular coupling, without requiring detailed information on experimentally inaccessible kinetic rates of branch retraction, soma relocation, and branch regrowth.

At each time step, we compute the center of mass (CoM) **r**_*i*_ and radius of gyration *R*_*i*_ for each cell *i* from its current branch coordinates using the minimum-image convention for the periodic boundary. To restrict interactions to nearby cells and avoid spurious long-range forces, we determine the neighbor set 𝒩_*i*_ for each cell via Delaunay triangulation of the CoM positions under periodic boundary conditions. The repulsive displacement of cell *i* is then given by

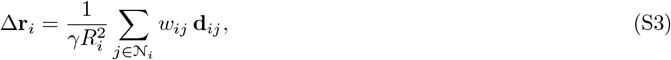

where **d**_*ij*_ = **r**_*i*_ −**r**_*j*_ is the minimum-image CoM separation vector, and 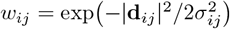 is a Gaussian soft-core weight with width 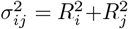. The Gaussian kernel ensures that the interaction force decays smoothly with distance and vanishes for well-separated cells, without introducing a hard cutoff. Following [79], we model the effective drag coefficient to scale as the square of the cell’s radius of gyration, 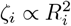, reflecting a contact-dominated friction driven by the number of adhesive contacts. The displacement Δ**r**_*i*_ is applied rigidly to all branch coordinates and the soma of cell *i* at the end of each time step, preserving the intrinsic morphology of the cell. This is a simplified assumption justified by the lack of experimental constraints on the microscopic branch retraction and regrowth kinetics, and provides a minimal effective intercellular repulsive coupling controlled by a single friction coefficient *γ*.

##### Density-dependent growth arrest rule

To model microglial growth arrest during development, we require a mechanistic rule that captures local spatial constraints without introducing unmeasured biological parameters (e.g., molecular diffusion coefficients or branch-retraction rates). To achieve this, we develop a coarse-grained, density-dependent growth arrest model based on local territorial packing.

Rather than simulating explicit branch-to-branch steric interactions, which are computationally expensive and highly sensitive to the microscopic details of the stochastic branching process, we define an “interaction zone” for each cell using a bounding disk. For any branched cell *i* at a given developmental timestep, we calculate its center of mass, **r**_cm,*i*_, based on the coordinates of all its constituent branches. We then determine the cell’s maximal reach, *R*_*i*,max_, defined as the distance from the center of mass to its furthest branch tip. The cell’s territorial footprint is thus approximated as a disk of area 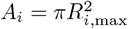, providing a proxy for the local area occupied by the growing branches of the cell.

We define two cells, *i* and *j*, as interacting neighbors if their bounding disks overlap, satisfying the condition |**r**_cm,*i*_ − **r**_cm,*j*_| *< R*_*i*,max_ + *R*_*j*,max_. For any active cell *c* with *M* identified neighbors, the total territorial area claimed by this local collective is

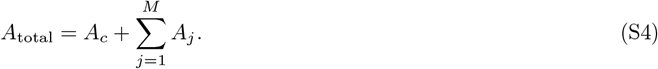

Growth arrest is triggered when this local collective packing reaches a fundamental geometric limit. We utilize the optimal packing fraction for uniform circles in two dimensions, defined by the hexagonal close-packing threshold 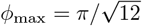. Given a global cellular density *ρ* = *N/A*_sim_ (where *N* is the total number of cells and *A*_sim_ is the total simulated tissue area), the expected available area for *M* + 1 cells is (*M* + 1)*/ρ*. We therefore arrest the branching process of cell *c* when the combined territorial area of the cell and its neighbors exceeds the theoretical packing limit of their shared space:

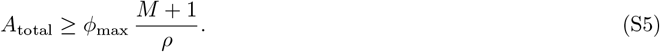

This local rule provides a parameter-free mechanism for density sensing, mapping local crowding directly onto the geometric constraints of 2D space-filling.

#### S1.3. Model sensitivity and parameter values

The collective BARW framework rests on two key mechanistic ingredients: intercellular repulsion and density-dependent growth arrest. Here we describe simulation results under different modeling assumptions to test the influence of these rules on cell growth and identify the conditions under which robust collective patterning occurs.

##### Role of individual mechanistic rules

We sequentially disabled the two collective mechanisms of repulsion and growth arrest to isolate their respective contributions, comparing three reduced models against the full control setup (Supp. Fig. **S2**).

i. *Without repulsion, without growth arrest* We initialized *N* cells with randomly placed soma and evolved them independently according to the single-cell BARW rules. In the absence of any collective interaction, cells grow without sensing their neighbors except through contact-mediated branch termination, with branches crossing freely into neighboring Voronoi territories. As a consequence, the fraction of branches remaining within each cell’s Voronoi domain decreased monotonically over time (Supp. Fig. **S2**b). Convex hull areas did not saturate but instead continued growing to approximately twice the values reached in the control at the same time point (Supp. Fig. **S2**c), and local tiling likewise failed to plateau (Supp. Fig. **S2**d). Since soma positions remained pinned to their initial random positions throughout, no regularity in soma positions could emerge, and hyperuniformity analysis is therefore not applicable.
ii. *With repulsion, without growth arrest* Here we activated the repulsion rule but disabled growth arrest. Cells could now reposition their soma in response to neighbors, and the resulting repulsive dynamics drove soma toward more regularly spaced configurations, producing hyperuniform-like positional order. However, without a growth arrest mechanism, branches continued to extend and invade neighboring territories. Territory crossing remained substantial at the final time point, with approximately 35% of branches crossing into neighboring Voronoi domains, compared to over 95% domain confinement in the full control (Supp. Fig. **S2**b). Convex hull areas and local tiling likewise behaved similarly to the no-repulsion case, as cells kept growing across domain boundaries (Supp. Fig. **S2**c-d).
iii. *Without repulsion, with growth arrest* We next activated growth arrest but disabled repulsion. Branch confinement within Voronoi domains was slightly reduced relative to the control (85–90%), as cells initialized in locally dense regions may cross the static Voronoi boundaries of their neighbors before growth arrest is triggered (Supp. Fig. **S2**b). The tiling dynamics were nevertheless qualitatively similar to the full control: convex hull areas and local tiling saturated at comparable values (*ϕ*≃ 0.5), confirming that growth arrest is the primary driver of territorial self-limitation (Supp. Fig. **S2**c-d). Critically, since soma positions remained fixed throughout, no positional ordering could develop: all positional statistics (Voronoi area distributions, Delaunay edge lengths, number fluctuations) remained frozen at their initial random values, and the scaling exponent of number fluctuations stayed at *λ* = 2 (Poisson) throughout, as discussed further below. Together, these three controls establish that territorial confinement requires growth arrest, while the approach to hyperuniform positional order requires repulsion: both ingredients are necessary to reproduce the full phenomenology observed in the control simulations and in experiments.

**Supplementary Figure S1.**
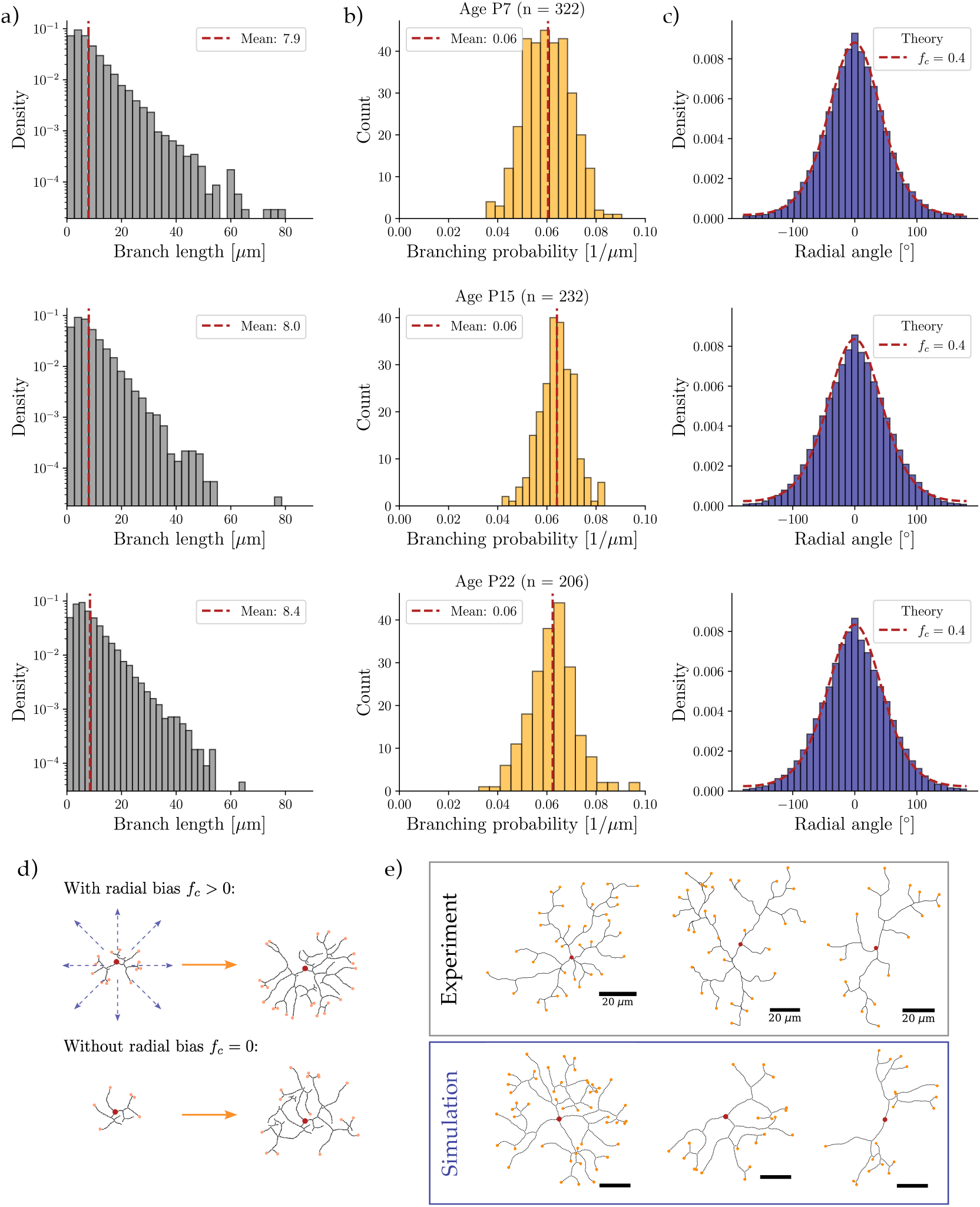
Statistical features of individual microglial cells. **a)** Branch length distributions show an exponential decay, as predicted by a Poisson branching process with fixed branching probability. Mean branch length (∼ 8 *µ*m) is conserved across development (top to bottom). **b)** Branching probability inferred from branch length distributions at different developmental ages (top to bottom). **c)** Branch orientations of individual microglia, measured by the radial angle relative to the soma position. The theoretical prediction from the radially guided BARW model (dashed line) indicates a non-zero guidance parameter *f*_*c*_. **d)** Illustration of the effect of radial bias on simulated single-cell morphologies. Top: BARW with radial guidance (*f*_*c*_ *>* 0). Bottom: unbiased BARW (*f*_*c*_ = 0). **e)** Representative single-cell morphologies from experiments (top) and simulations (bottom) using experimentally inferred parameters. Terminal branches and somata are indicated by orange and red dots, respectively.

**Supplementary Figure S2.**
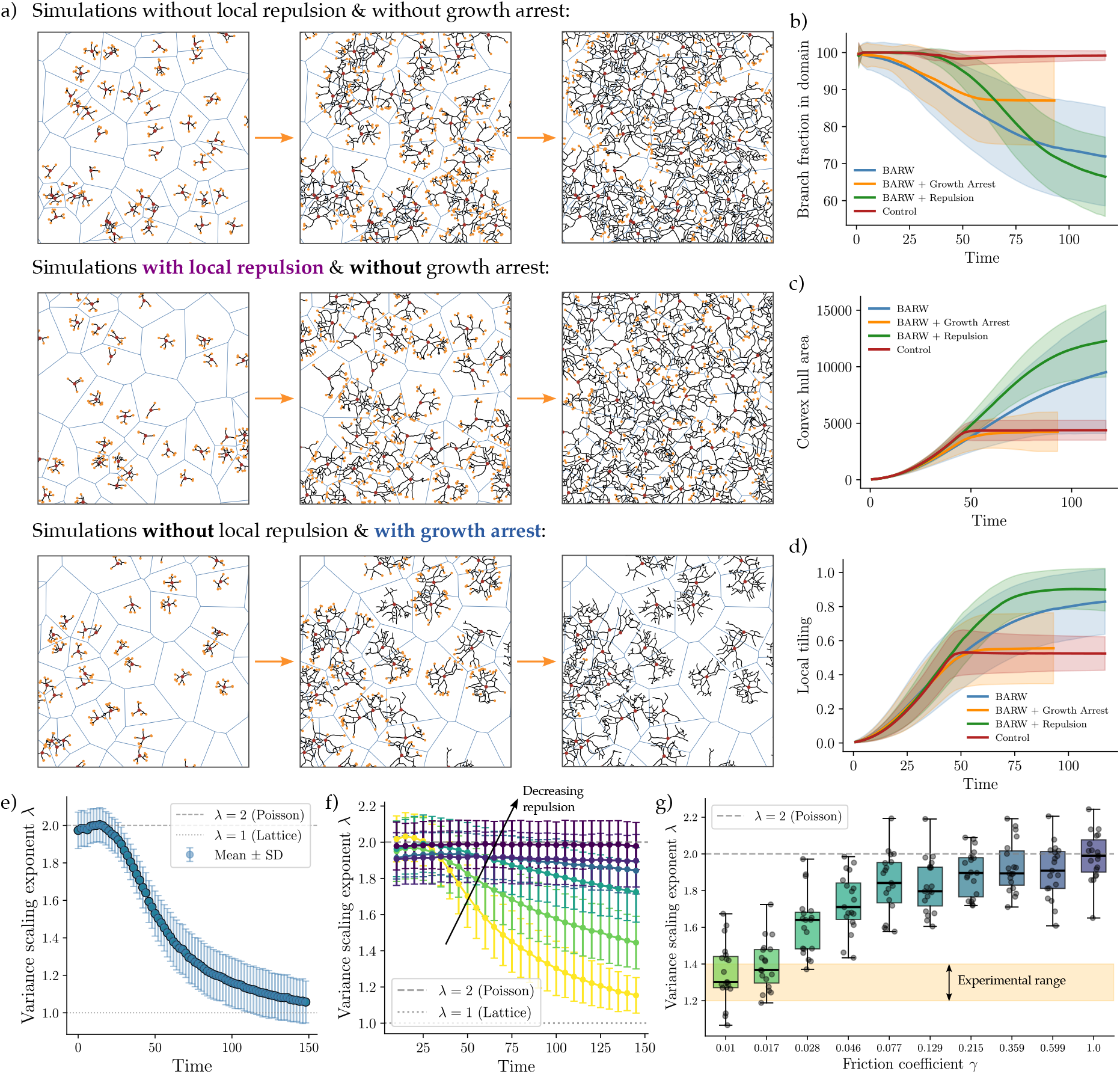
Simulations of *N* -cell BARWs under different modeling assumptions. **a)** Collective branching patterns in BARW simulations without repulsion and without growth arrest (top), with repulsion and without growth arrest (middle), and without repulsion and with growth arrest (bottom). Time evolution proceeds in the direction of the arrows. **b)** Fraction of each cell’s branch segments lying within its own Voronoi domain, for different model assumptions. **c)** Convex hull area evolution for different model assumptions. **d)** Local tiling metric evolution for different model assumptions. In **b)**–**d)**, solid lines and shaded regions denote mean ±SD. **e)** Time evolution of the variance scaling exponent in the control simulations. Dashed line: Poisson exponent *λ* = 2. Dotted line: lattice exponent *λ* = 1. Markers and error bars: mean ±SD from *n* = 50 independent simulations. **f)** Time evolution of the variance scaling exponent for varying repulsion strength, controlled by the friction coefficient *γ* (light to dark: increasing *γ*, i.e. decreasing mobility). Markers and error bars: mean ±SD from *n* = 20 simulations. **g)** Distribution of the variance scaling exponent *λ* for different values of the friction coefficient *γ* at a time point matched to late-stage experiments. The late-stage experimental range *λ* = 1.2–1.4 is highlighted by the horizontal shaded band (orange). Box plots show the median (center line), interquartile range (box), whiskers (1.5 × IQR), and individual mean exponents (points) from *n* = 20 simulation repeats.

##### Robustness to cell density

To test whether the proposed mechanism generalizes beyond the specific cell number used in the control simulations, we varied the number of cells *N* systematically while keeping the domain side length fixed at *L* = 1000, thereby spanning a wide range of initial cell densities *ρ*_0_ = *N/L*^2^. Across all densities, local tiling increased monotonically during growth and saturated at *ϕ* ≃0.5–0.6, consistent with the control and experimental values, although higher-density conditions reached saturation faster due to earlier onset of the growth arrest criterion (Supp. Fig. **S5**e). The hyperuniformity scaling exponent followed a qualitatively similar trajectory at all densities, decreasing from *λ* = 2 at early times toward *λ <* 1.2 at the final simulated time (Supp. Fig. **S5**d). These results confirm that the mechanism robustly generates hyperuniform tiling across a wide range of cell densities, thereby validating that the *rd10* phenotype with impaired tiling and fluctuation suppression cannot be a direct consequence of increased cell density. This analysis also validated the *A*_*c*_ ≃ 0.5 *ρ*^−1^ scaling of individual cell territory area observed in the experimental data (Fig. **S3**b).

**Supplementary Figure S3.**
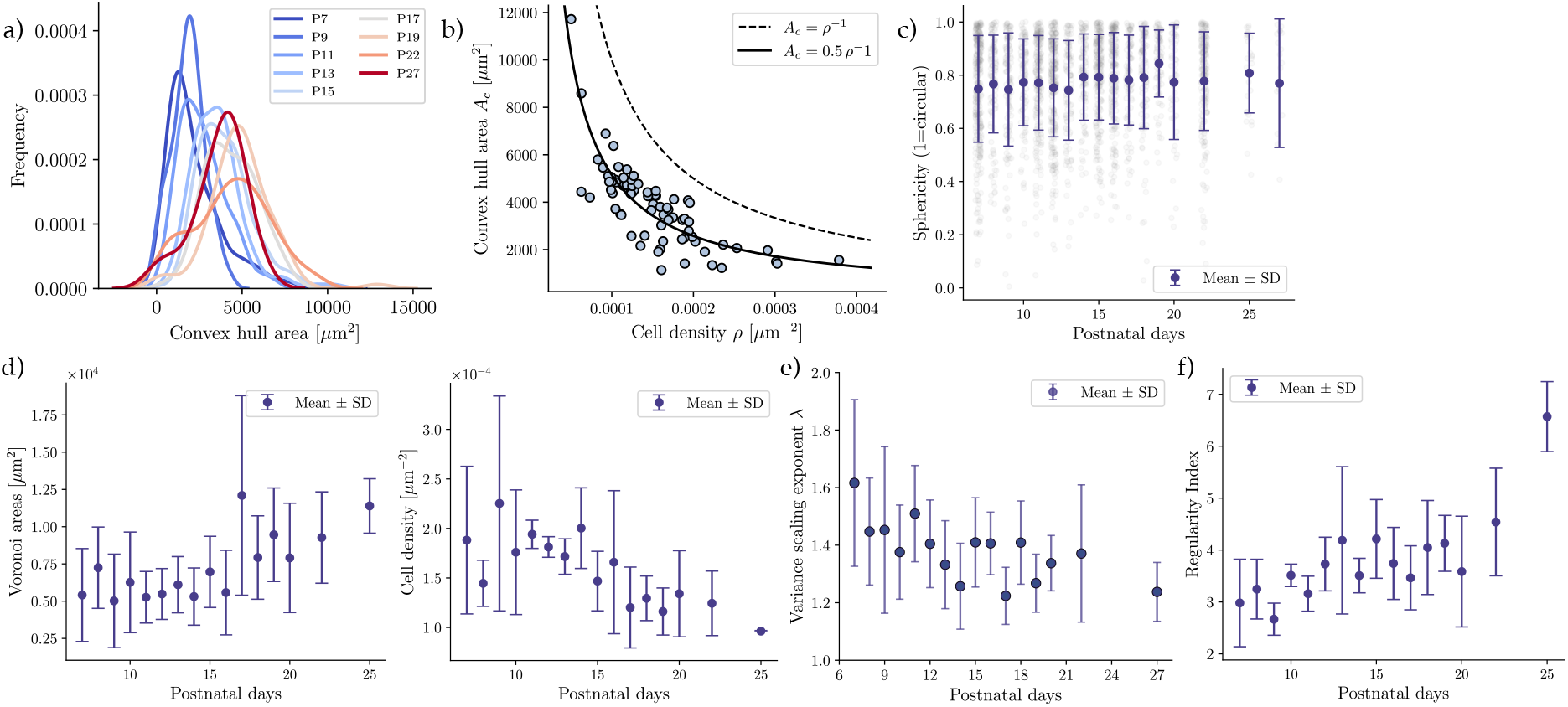
Experimental statistics on microglial growth. **a)** Distributions of convex hull areas across development (color-coded). **b)** Convex hull area as a function of cell density for each sample at all available developmental time points. Dashed and solid lines show the scaling relationships *A*_*c*_ = *ρ*^*−*1^ and *A*_*c*_ = 0.5 *ρ*^*−*1^, respectively. **c)** Sphericity metric across developmental time points. **d)** Evolution of Voronoi area (left) and cell density (right) across developmental time points. **e)** Evolution of variance scaling exponent *λ* across developmental time points. **f)** Voronoi regularity index across developmental time points. In **c)**–**f)**, markers and error bars indicate mean ± SD.

**Supplementary Figure S4.**
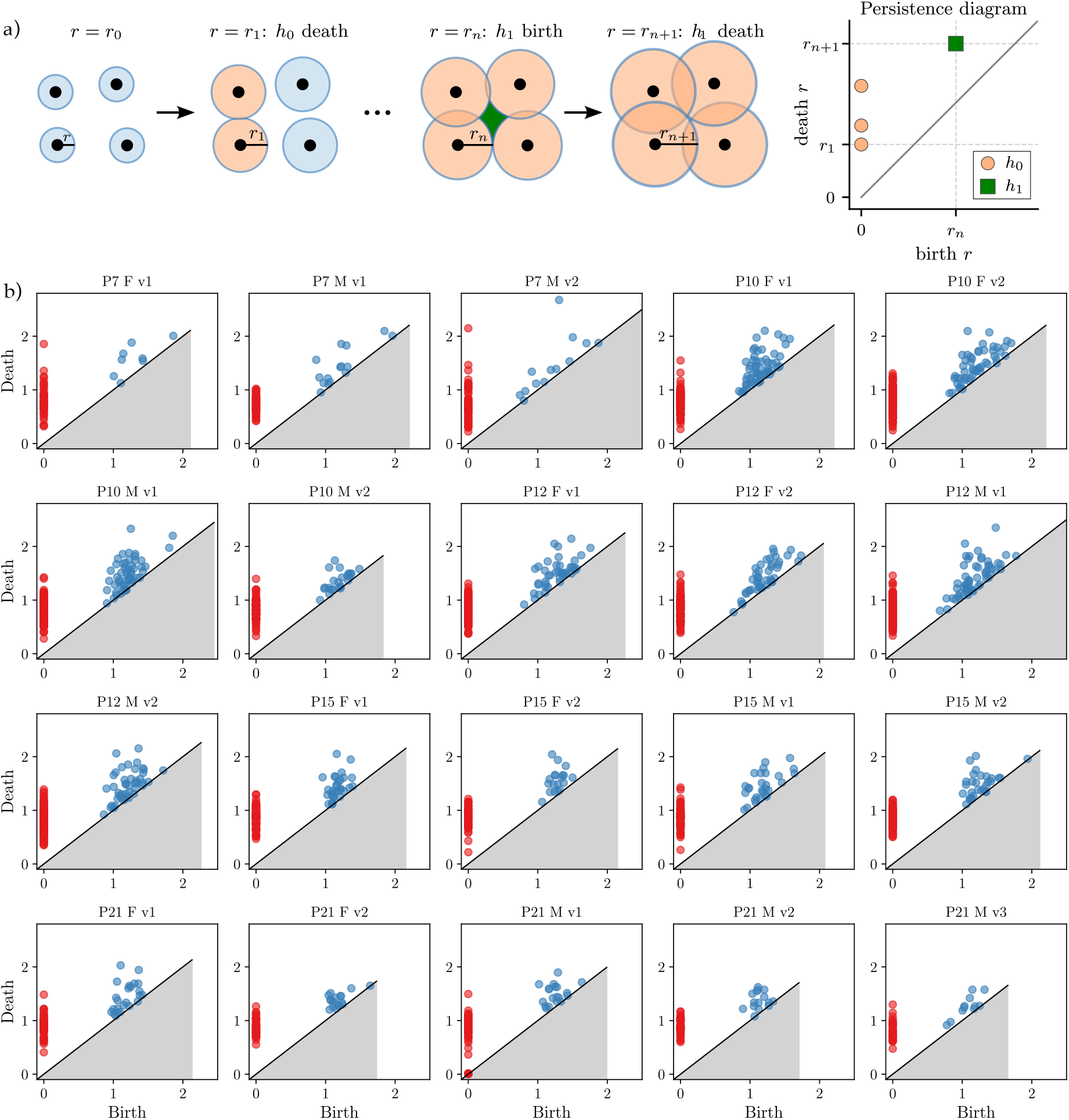
Full TDA analysis on experimental soma coordinates. **a)** Schematic illustration of persistent homology on a 2D point pattern. Disks of increasing radius *r* are grown around each point, with the radius shown for one disk per panel. At *r* = *r*_1_, two disks first touch, forming an edge and marking the death of a connected component (*h*_0_). At *r* = *r*_*n*_, the overlapping disks enclose a gap, marking the birth of a topological hole (*h*_1_, green). At *r* = *r*_*n*+1_, the hole is filled as disks continue to grow, marking its death. The persistence diagram (right) summarizes each feature with marked birth and death radii for *h*_0_ (disk) and *h*_1_ (square) features. **b)** Vietoris-Rips persistence diagrams for soma coordinate point clouds at different developmental time points. Connected components *h*_0_ (dimension 0) are shown in red, holes *h*_1_ (dimension 1) in blue. Plot titles indicate postnatal day (PX), sex (F/M), and sample index (vX).

**Supplementary Figure S5.**
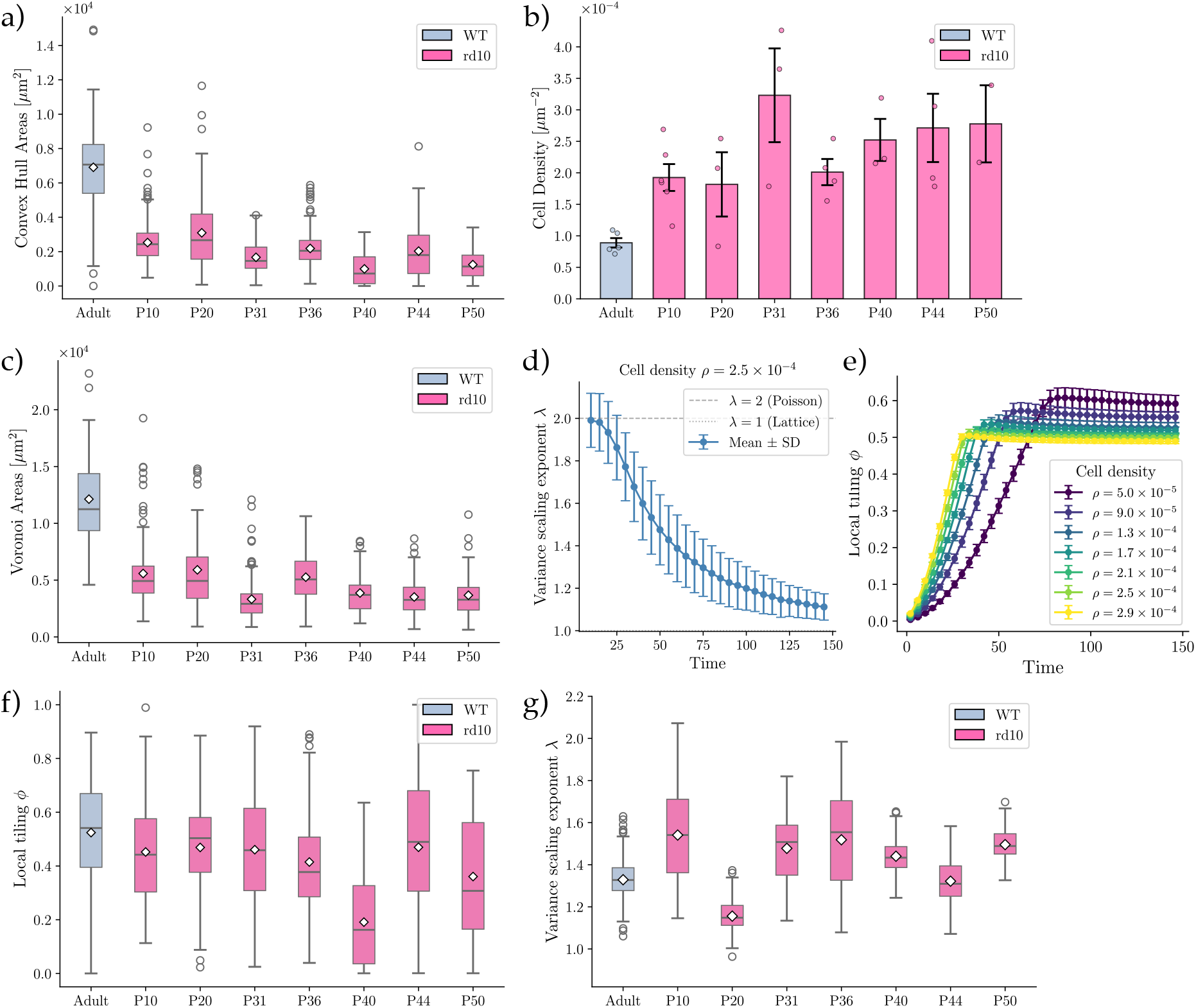
Statistics on the *rd10* phenotype and related simulation results. **a)** Box plot of convex hull areas across developmental time points. **b)** Mean cell density across developmental time points. Error bars: SD. Points: individual values per age. **c)** Box plot of Voronoi areas across developmental time points. **d)** Variance scaling exponent over time from stochastic branching simulations with WT parameters but cell density matched to the late-stage *rd10* phenotype. Dashed line: Poisson exponent *λ* = 2. **e)** Time evolution of the local tiling metric *ϕ* from stochastic branching simulations with WT parameters across different cell densities *ρ* (color-coded). In **d)**–**e)**, markers and error bars show mean ±SD from *n* = 20 simulation repeats. **f)** Box plot of local tiling *ϕ* across developmental time points. **g)** Box plot of variance scaling exponent *λ* across developmental time points. In **a)**–**c)** and **f)**–**g)**, *rd10* data cover time points from P10 to P50 (pink), compared with the WT adult dataset (gray). Box plots show median (center line), mean (center marker), interquartile range (box), whiskers (1.5× IQR), and outliers (individual points).

##### Sensitivity to repulsion strength

The friction coefficient *γ*, which controls the magnitude of soma displacement due to repulsive interactions (Eq. S3), is not directly constrained by experimental data and constitutes the main free parameter of the collective model. Its role can be understood physically as setting the timescale of soma repositioning relative to the timescale of branch growth: if *γ* is too large, mechanical relaxation is slow, and growth arrest is triggered before cells have had time to reach regularly spaced configurations, suppressing positional ordering. Conversely, for sufficiently small *γ*, cells reposition rapidly during growth and the system evolves toward the hyperuniform state observed in the control.

To quantify this dependence and constrain the choice of *γ*, we performed additional simulations spanning two orders of magnitude around the control value *γ* = 0.01 (see Table S1). We found that as *γ* increases, the decay of the number fluctuation scaling exponent from the Poisson value *λ* = 2 slowed progressively, with the large-*γ* limit reproducing the behavior of the growth-arrest-only model, indicating that no positional regularity emerged over time (Supp. Fig. **S2**f). We then examined the distribution of *λ* values at simulation time points matched to the late experimental stage P30, and compared the scaling exponents with the range *λ* = 1.2–1.4 observed in latestage experiments (Supp. Fig. **S3**e). Scaling exponents from the *γ* = 0.01 simulations clearly lay within this range across independent repeats, while *γ >* 0.02 already deviated strongly from the experimentally observed scaling regime (Supp. Fig. **S2**g). This comparison quantitatively constrained the value of *γ* used in the control simulations. Finally, local tiling dynamics remained relatively insensitive to *γ*, since growth arrest is governed primarily by the branching dynamics and not by soma displacement, consistent with the results of the growth-arrest-only control above.

##### Simulation parameters

All parameters used in the control collective simulations are listed in Table S1.

**TABLE S1.** Parameters used in the control *N* -cell BARW simulations.

| Parameter | Symbol | Value | Description |
| --- | --- | --- | --- |
| Territory size | $L$ | $1000 \ell$ | Side length of the periodic simulation box |
| Cell number | $N$ | 120 | Number of cells |
| Branching probability | $p_b$ | 0.06 | Per-step branching probability |
| Elongation angle | $\psi_e$ | $\pi/10$ | Maximum angular deviation per elongation step |
| Annihilation radius | $r_{\text{ann}}$ | $3 \ell$ | Sensing distance for contact-driven tip termination |
| Radial guidance strength | $f_c$ | 0.4 | Strength of outward radial guidance cue |
| Friction coefficient | $\gamma$ | 0.01 | Inverse mobility for cell repulsion |

### Supplementary Section S2: Persistent homology analysis

Persistent homology identifies patterns in data by applying a filtration that tracks topological features across multiple scales. A commonly used filtration function is the height function [80]. In the context of point clouds, distancebased filtrations such as the Vietoris–Rips and Čech filtrations are widely employed to construct a growing family of simplicial complexes from a set of points [80, 81].

In particular, the Vietoris–Rips filtration, which is often preferred for its computational efficiency [82], is built by connecting pairs of points whose distance is at most a scale parameter *r*, and including a *k*-simplex whenever all its vertices are pairwise connected. For example, a 1-simplex represents an edge between two points at distance at most *r*, contributing to the formation of connected components. The *h*_0_ homology group counts the connected components, which correspond to distinct clusters in the data. A 2-simplex corresponds to three pairwise-connected points forming a filled triangle. The *h*_1_ homology group captures one-dimensional holes or loops, representing cycles that are not filled in by higher-dimensional simplices.

As the scale parameter *r* increases, this construction yields a nested sequence of simplicial complexes, studying how topological features, such as connected components and loops, appear and disappear, thereby revealing the underlying structure of the metric space.

### Supplementary Section S3: Langevin model for growing soft disks

Here, to complement the microscopic BARW simulations and study the mechanism of hyperuniform tiling driven by the coupled dynamics of repulsion and growth arrest at a coarser level of description, we represent each cell as a soft, growing disk. Our motivation is to isolate the role of mechanical cell-cell interactions and contact-inhibited growth in driving spatial order, independently of the detailed branching morphology. This approach follows the Langevin framework previously applied to growing bacterial colonies [60] interacting via a Hertzian elastic contact potential, adapted here to isotropically growing disks under overdamped dynamics.

#### Dimensional equations of motion

We consider *N* disks in a two-dimensional periodic domain of side length *L*, each characterized by a position **r**_*i*_(*t*) and a radius *R*_*i*_(*t*). Cell positions follow overdamped dynamics

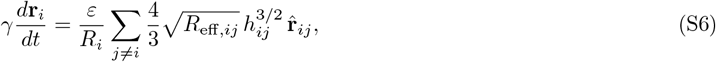

where *γ* is a drag coefficient, *ε* is an elastic modulus setting the repulsion strength, 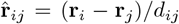 is the unit vector along the line of disk centers, *h*_*ij*_ = max(*R*_*i*_ + *R*_*j*_ − *d*_*ij*_, 0) is the pairwise overlap, and *R*_eff,*ij*_ = *R*_*i*_*R*_*j*_*/*(*R*_*i*_ + *R*_*j*_) is the effective contact radius. The 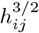 dependence follows from Hertz contact theory, where the elastic restoring force between two compressed bodies scales as the overlap to the power 3*/*2. The cell radius evolves as

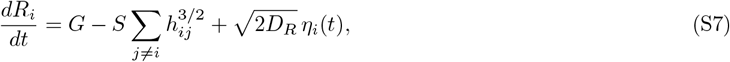

where *G* is the intrinsic growth speed, *S* is the rate of contact-inhibited shrinkage per unit overlap, *D*_*R*_ is the radial diffusion coefficient, and *η*_*i*_(*t*) are independent Gaussian white noises with zero mean and unit variance. Growth is thus suppressed by mechanical contact, with net radial expansion only when the shrinkage speed 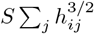 is smaller than *G*.

#### Nondimensionalization

Rescaling lengths by the initial radius *R*_0_ and time by the growth timescale *T*_0_ = *R*_0_*/G* yields the dimensionless form presented in the main text, with control parameters

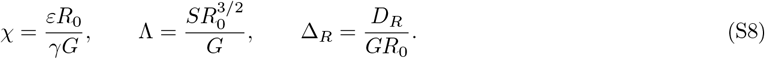

The parameter *χ* quantifies the strength of mechanical repulsion relative to growth, Λ sets the competition between contact-induced shrinkage and intrinsic growth (with Λ *>* 1 implying net shrinkage under crowding), and Δ_*R*_ is kept small throughout (Δ_*R*_ = 10^−4^) so that stochasticity enters only as a weak perturbation.

#### Implementation

All pairwise distances are computed under the minimum-image convention for the periodic domain. The equations of motion are integrated with a forward Euler scheme at fixed time step *dt* = 0.05 *T*_0_ until *T* = 100. Radii are prevented from becoming negative. The behavior of the system as a function of Λ and *χ* is characterized via systematic parameter sweeps, with *n* = 20 independent realizations per parameter combination. The final-state configuration of each replicate is then characterized using the following metrics.

#### Packing fraction

Naively estimating the packing fraction as 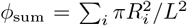 overestimates area coverage when disks overlap. To correct for this, we computed the true packing fraction *ϕ* as the union area of all disks normalized by *L*^2^. Each disk was approximated as a 64-sided polygon, and image copies were generated on the opposing side of the domain for any disk whose boundary crossed the simulation boundary. The union *A*_union_ of all primary disks and their periodic images was computed using the shapely library, and the result was intersected with the simulation box [0, *L*]^2^ to obtain the covered area within the domain.

#### Tiling efficiency

To quantify how efficiently the cells collectively tile the available space without overlap, we define the overlap cost

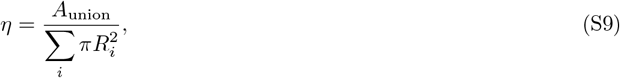

where *A*_union_ is the true union area computed above. A value of *η* = 1 indicates no disk-disk overlap, while *η <* 1 reflects the fraction of total disk area lost to overlaps. The combined tiling efficiency metric reported in the main text is then

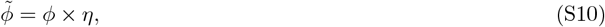

which simultaneously penalizes poor coverage and excessive overlap, and is maximized by configurations that tile the plane efficiently with minimal disk-disk overlaps.

#### Number fluctuation exponent

To assess the degree of spatial regularity in disk center positions, we measured the variance 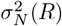 of the number of cell centers falling within circular observation windows of radius *R*, placed at *n* = 50 random positions within the domain, for a set of logarithmically spaced radii *R*∈ [*R*_min_, *R*_max_], following the same analysis pipeline for simulation and experimental data (see below). We extracted the exponent *λ* by a linear fit to log 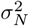 versus log *R*. To obtain a robust estimate, the window placement was randomized and the variance scaling exponent calculation was repeated *M* = 50 times per configuration.

### Supplementary Section S4: Data analysis

#### S4.1. Individual cell statistics

Morphological reconstructions of individual microglia were obtained as SWC-format files as described in the Methods section of the main text. Each SWC file encodes the branched cell as a directed tree, where every node carries a unique identifier, coordinate values, and a pointer to its parent node. The root node (parent identifier −1) corresponds to the soma. We parsed the SWC tree into a set of branch segments, defined as paths between consecutive branch points, or between a branch point and a terminal leaf node. To extract branching statistics, we traversed the tree from each leaf toward the root, collecting nodes until a branch point or the soma was reached. The Euclidean length of each segment was computed by summing the distances between consecutive nodes along the path.

##### Branch length distributions

The lengths of all branch segments, recorded across all reconstructed cells, provided a characterization of the typical branching scale of individual cells. More importantly, an exponentially distributed branch length indicates an underlying Poisson branching process, as we observed (Supp. Fig. **S1**a), providing a quantitative validation of the stochastic BARW framework.

##### Branching probability

For each root-to-leaf lineage, we estimated the branching probability per unit length as

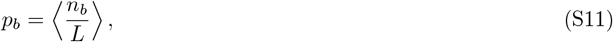

where *n*_*b*_ is the number of branch points encountered along the lineage, *L* is its total path length in units of 𝓁, and the average is taken over all lineages and all reconstructed cells.

##### Radial guidance strength

To capture local branch orientation at finer resolution than the full orientation of an entire branch segment, we subdivided each branch segment into smaller blocks of a fixed number of SWC edges. For each block, we computed two vectors in the 2D plane: the block direction *φ*, given by the net displacement from the block start to its end, and the local radial direction *θ*, pointing from the soma to the block midpoint. The signed radial angle *ψ* = *φ*− *θ* is wrapped to [− *π, π*], so that *ψ* = 0 corresponds to a block pointing directly away from the soma, *ψ* = ± *π/*2 to a tangentially oriented block, and *ψ* = ±*π* to a block directed back toward the soma. These angular variables map directly onto the variables of the radially guided BARW model described above. Pooling *ψ* across all blocks and all reconstructed cells yields an angular distribution reflecting the degree of radial bias. Theoretically, BARW under guidance predicts branch orientations to follow a von Mises distribution [32]

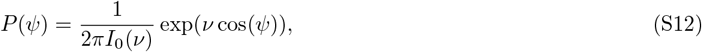

where *I*_0_(*ν*) is the modified Bessel function of the first kind of order zero and the concentration parameter *ν* is given by

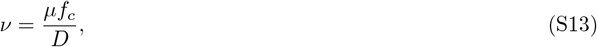

where

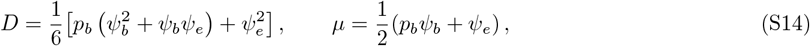

with *ψ*_*b*_ the maximum branching angle and *ψ*_*e*_ the maximum elongation angle. Fitting the experimentally obtained radial angle distribution to Eq. S12 thus provided an estimate of the radial guidance strength *f*_*c*_, confirming the presence of a guided BARW process in individual microglia (Supp. Fig. **S1**c).

#### S4.2. Collective spatial statistics

We applied the following metrics to soma coordinate data extracted from both experimental retinal subregions and simulations, and explicitly note where the experimental and simulation implementations differ in boundary handling.

##### Voronoi tessellation and boundary handling

All spatial statistics involving territorial assignments are built on a Voronoi tessellation of soma positions. Because Voronoi cells near the boundary of the field of view are artificially large or unbounded, a consistent boundary correction is applied in both cases before any metric is computed.

For experimental data, we used the global convex hull of all branch coordinates across all cells in the field of view as the boundary polygon. Each Voronoi cell is clipped to this hull using polygon intersection (with the Shapely library), and only cells whose clipped polygon is valid and non-degenerate are retained for analysis.

For simulations, the domain is periodic and the Voronoi tessellation is computed on a 3×3 mirror-tiled copy of the soma coordinates (covering all 9 periodic images), which correctly handles cells near the domain boundary. Voronoi cells whose vertices fall outside the simulation box [0, *L*]^2^ are excluded from metric calculations.

##### Convex hull area and local tiling metric

To quantify the spatial territory occupied by each cell, we computed the convex hull of its branch coordinates. The convex hull area *A*_*c*_ provides a coarse-grained estimate of the cell’s territorial footprint. The local tiling fraction *ϕ*_*c*_ is defined for each cell *c* as the ratio of the intersection area between the cell’s convex hull polygon and its corresponding (clipped) Voronoi domain to the Voronoi domain area:

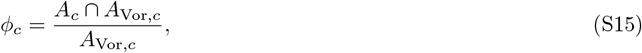

where *A*_Vor,*c*_ denotes the area of the (clipped) Voronoi domain of cell *c*. A value of *ϕ*_*c*_ = 1 indicates that the cell’s convex hull exactly fills its Voronoi territory, while *ϕ*_*c*_ *<* 1 reflects either incomplete coverage or hull-Voronoi mismatch due to irregular cell shape. We tracked the mean local tiling fraction ⟨*ϕ*⟩ across all valid interior cells as a function of developmental time.

##### Branch confinement within Voronoi domains

As a more direct measure of territorial overlap between neighboring cells, we computed the fraction of branch coordinates of each cell that fall within its own Voronoi domain. For each cell *c*, all branch coordinate points are assigned to their nearest soma using a *k*-d tree query on soma positions. The branch confinement fraction is then

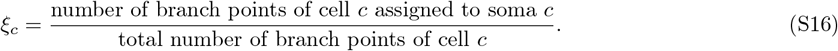

A value of *ξ*_*c*_ = 1 indicates complete territorial confinement, while *ξ*_*c*_ *<* 1 reflects branches crossing into a neighboring cell’s Voronoi domain. We tracked the mean confinement fraction ⟨*ξ*⟩ across all valid cells over time.

##### Edge lengths and domain regularity

To characterize the regularity of soma spacing, we computed two complementary statistics from the Voronoi tessellation and Delaunay triangulation of soma positions. First, we calculated edge lengths as the distances between all pairs of Delaunay-connected somas. Second, we quantified the regularity of Voronoi domains via

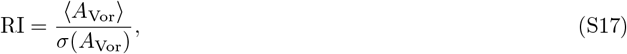

where ⟨*A*_Vor_⟩ is the mean and *σ*(*A*_Vor_) the standard deviation of the Voronoi area distribution. Higher RI values indicate more uniform Voronoi domains.

##### Number variance and hyperuniformity exponent

To quantify long-range density fluctuations in soma positions, we measured the number variance 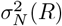: the variance of the count of soma centers within circular observation windows of radius *R*, placed at *n* = 50 random positions within the domain. This was computed at logarithmically spaced values of *R* between *R*_min_ and *R*_max_, chosen to span more than one order of magnitude within the available field of view, while omitting values *R > L/*6 to avoid repeated counts over overlapping windows and artificially reduced exponents. For a Poisson (uncorrelated) point pattern, 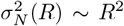; for a perfect lattice, 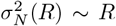; and for a hyperuniform system, 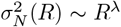 with *λ <* 2. We extracted the scaling exponent *λ* by a linear fit to log *σ*^2^ versus log *R*. To obtain robust estimates, window placement was randomized and the slope calculation was repeated *N*_*t*_ = 50 times per time point and per simulation replicate, and we reported the mean and standard deviation of *λ* across these trials.

The implementation is identical for experiments and simulations, with the observation domain set to the bounding box of the soma coordinates in the experimental case and to [0, *L*]^2^ in the simulated case.

##### Structure factor

The isotropic structure factor *S*(*k*) is computed from soma positions as

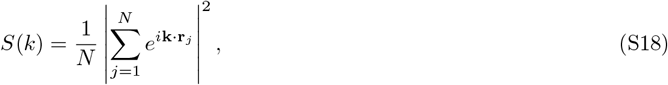

evaluated on the discrete set of wavevectors **k** compatible with the domain geometry, and then radially averaged into bins of width Δ*k* to yield the isotropic *S*(*k*). Wavevectors for which fewer than *n*_min_ = 3 full wavelengths fit within the box are discarded to avoid finite-size artefacts at small *k*. The resulting *S*(*k*) curves were averaged across simulation replicates at each time point and compared directly to the experimental *S*(*k*) obtained from the same pipeline applied to experimental soma coordinates.

### S4.3. Implementation details for persistent homology

Persistent homology computations were carried out using the GUDHI library (https://gudhi.inria.fr/). For each point cloud of soma coordinates (experimental field of view or simulation box), we constructed the Vietoris-Rips filtration up to maximum simplex dimension 2, yielding *h*_0_ (connected components) and *h*_1_ (holes) persistence diagrams, as described in Section S2. Point clouds with fewer than 40 soma were excluded from the analysis.

#### Coordinate normalization

To enable direct comparison between experimental fields of view and simulations, which differ in absolute spatial scale and cell density, all soma coordinates were normalized before computing persistence. For a point cloud of *N* somas within a domain of area *A*, coordinates were rescaled by the mean inter-soma spacing

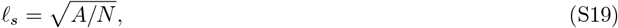

so that the normalized coordinates are dimensionless and the typical inter-soma distance is of order unity. This normalization ensures that persistence diagram features reflect the topological structure of the pattern independently of absolute scale or density.

#### Feature extraction

From each persistence diagram, *h*_0_ and *h*_1_ features were extracted as follows. For *h*_0_, each feature is born at filtration value 0 (all points start as isolated components) and dies at the scale at which its corresponding connected component merges with another. The *h*_0_ lifetime is *τ*_0_ = *r*_death_, and the single feature with infinite death value (the last remaining connected component) is excluded. For *h*_1_, each feature is born at the scale *r*_birth_ at which a loop first forms and dies at the scale *r*_death_ at which it is filled in. The *h*_1_ lifetime is *τ*_1_ = *r*_death_ −*r*_birth_, and only finite features are retained. In addition to lifetime, we computed the *h*_1_ characteristic scale

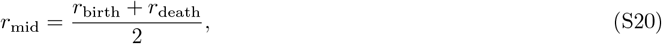

which characterizes the spatial scale at which a loop is most persistent and provides a complementary descriptor to the lifetime.

#### Pooling and comparison

For simulations, persistence diagrams were computed at selected time points corresponding to the developmental stages P7, P12, and P21 in experiments, and pooled across 50 independent simulation replicates, yielding distributions of *h*_0_ and *h*_1_ features at each time point. For experimental data, diagrams were computed for each field of view independently and pooled across all available fields at the corresponding developmental ages. The resulting lifetime and characteristic scale distributions from simulations and experiments were then compared as normalized histograms.

